# The costs of competition: high social status males experience accelerated epigenetic aging in wild baboons

**DOI:** 10.1101/2020.02.22.961052

**Authors:** Jordan A. Anderson, Rachel A. Johnston, Amanda J. Lea, Fernando A. Campos, Tawni N. Voyles, Mercy Y. Akinyi, Susan C. Alberts, Elizabeth A. Archie, Jenny Tung

## Abstract

Aging, for virtually all life, is inescapable. However, within populations, biological aging rates vary. Understanding sources of variation in this process is central to understanding the biodemography of natural populations. We constructed a DNA methylation-based age predictor for an intensively studied wild baboon population in Kenya. Consistent with findings in humans, the resulting “epigenetic clock” closely tracks chronological age, but individuals are predicted to be somewhat older or younger than their known ages. Surprisingly, these deviations are not explained by the strongest predictors of lifespan in this population, early adversity and social integration. Instead, they are best predicted by male dominance rank: high-ranking males are predicted to be older than their true ages, and epigenetic age tracks changes in rank over time. Our results argue that achieving high rank for male baboons—the best predictor of reproductive success—imposes costs consistent with a “live fast, die young” life history strategy.

## Introduction

Aging, the nearly ubiquitous functional decline experienced by organisms over time^1^, is a fundamental component of most animal life histories^2^. At a physiological level, age affects individual quality, which in turn affects the ability to compete for mates and other resources, invest in reproduction, and maintain somatic integrity. At a demographic level, age is often one of the strongest predictors of survival and mortality risk, which are major determinants of Darwinian fitness. In order for patterns of aging to evolve, individuals must vary in their rates of biological aging. Thus, characterizing variation in biological aging rates and its sources—beyond simply chronological age—is an important goal in evolutionary ecology, with the potential to offer key insight into the trade-offs that shape individual life history strategies^3^.

Recent work suggests that DNA methylation data can provide exceptionally accurate estimates of chronological age^4^. These approaches typically use supervised machine learning methods that draw on methylation data from several hundred CpG sites, identified from hundreds of thousands of possible sites, to produce a single chronological age prediction^5–7^. Intriguingly, some versions of these clocks also predict disease risk and mortality, suggesting that they capture aspects of biological aging that are not captured by chronological age alone^8^. For example, in humans, individuals predicted to be older than their true chronological age are at higher risk of Alzheimer’s disease^9^, cognitive decline^9,10^, and obesity^11^. Accelerated epigenetic age is in turn predicted by environmental factors with known links to health and lifespan, including childhood social adversity^12,13^ and cumulative lifetime stress^14^. These observations generalize to other animals. Dietary restriction, for instance, decelerates biological aging based on DNA methylation clocks developed for laboratory mice and captive rhesus macaques, and genetic knockout mice with extended lifespans also appear epigenetically young for age^15–17^. However, while DNA methylation data have been used to estimate the age structure of wild populations (where birthdates are frequently unknown)^18–21^, they have not been applied to investigating sources of variance in biological aging in the wild.

To do so here, we coupled genome-wide data on DNA methylation levels with detailed behavioral and life history data available for one of the most intensively studied wild mammal populations in the world, the baboons of the Amboseli ecosystem of Kenya^22^. First, we calibrated a DNA methylation-based “epigenetic clock” and assessed the clock’s composition. Second, we compared the accuracy of this clock against other age-associated traits and between sexes. Third, we tested whether variance in biological aging was explained by socioenvironmental predictors known to impact fertility or survival in this population. Finally, we investigated an intriguing association between epigenetic age acceleration and male dominance rank. Our results show that predictors of lifespan can be decoupled from rates of epigenetic aging. However, other factors— particularly male dominance rank—are strong predictors of epigenetic clock-based age acceleration. These results establish the first epigenetic clock available for any wild nonhuman primate, and are the first to establish a link between social factors and epigenetic aging in any natural animal population. Together, they highlight potential sex-specific trade-offs that may maximize fitness, but also compromise physiological condition and potentially shorten male lifespan.

## Results

### Epigenetic clock calibration and composition

We used a combination of previously published^23^ and newly generated reduced-representation bisulfite sequencing (RRBS) data from 245 wild baboons (N = 277 blood samples) living in the Amboseli ecosystem of Kenya^22^ to generate a DNA methylation-based age predictor (an “epigenetic clock”^5,6^). Starting with a data set of methylation levels for 458,504 CpG sites genome-wide (Supplementary Figure 1; Supplementary Table 1), we used elastic net regression to identify a set of 573 CpG sites that together accurately predict baboon age to within a median absolute difference (MAD) of 1.1 years ± 1.9 s.d. (Figure 1; Supplementary Table 2; Pearson’s *r* = 0.762, p = 9.70 x 10^-54^; median adult life expectancy in this population is 10.3 years for females and 7.9 for males^24^). The choice of these sites reflects a balance between increasing predictive accuracy within the sample and minimizing generalization error to unobserved samples, using a similar approach as that used to develop epigenetic clocks in humans^5,6^ (see also Methods and Supplementary Figure 2).

**Figure 1.**
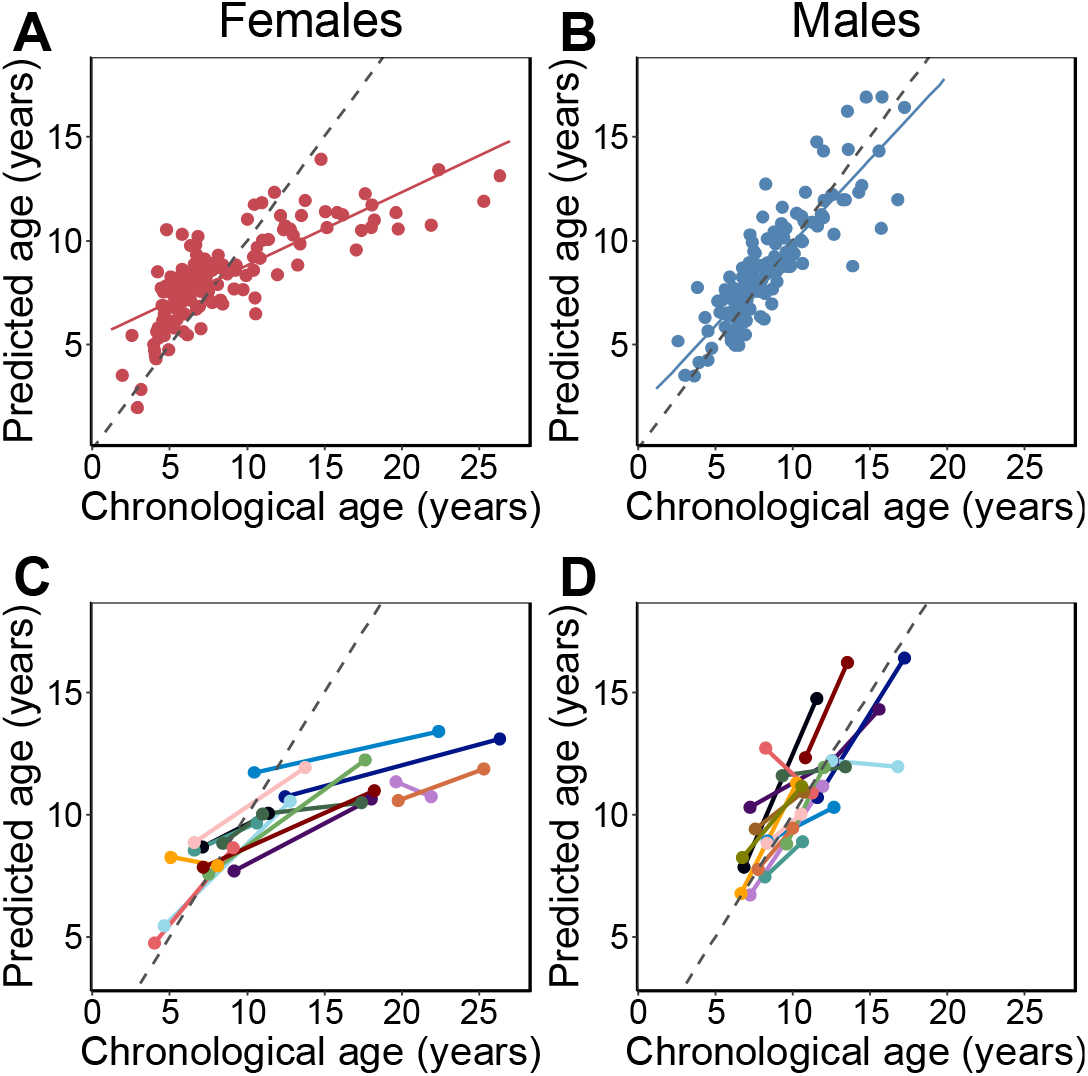
Epigenetic clock age predictions in the Amboseli baboons. Predicted ages are shown relative to true chronological ages for **(A)** females (Pearson’s *r* = 0.78, p = 6.78 x 10^-30^, N = 142 samples) and **(B)** males (*r* = 0.86, p = 5.49 x 10^-41^, N = 135 samples). Solid lines represent the best fit line; dashed lines show the line for y = x. **(C)** and **(D)** show predictions for individuals with at least two samples in the data set (N = 30; 14 females and 16 males). In 26 of 30 cases (87%), samples collected later were correctly predicted to be from an older animal.

Consistent with findings in humans^6^, clock sites are enriched in genes, CpG islands, promoter regions, and putative enhancers, compared to the background set of all sites we initially considered (Supplementary Figure 3; Fisher’s exact tests, all p < 0.05). Clock sites are also more common in age-associated differentially methylated regions in baboons (Supplementary Figure 3; sites that increase with age: log2[OR] = 4.189, p = 3.64 x 10^-9^; sites that decrease with age: log_2_[OR] = 5.344, p = 1.54 x 10^-8^)^25^, such that many, but not all, of the clock sites also exhibit individual associations between DNA methylation levels and age (Supplementary Figures 4 and 5; Supplementary Table 3). Additionally, clock sites were more likely to be found in regions that exhibit enhancer-like activity in a massively parallel reporter assay (sites that increase with age: log_2_[OR] = 2.685, p = 1.22 x 10^-2^; sites that decrease with age: log_2_[OR] = 4.789, p = 1.78 x 10^-5^)^26^ and in regions implicated in the gene expression response to bacteria in the Amboseli baboon population (overlap of lipopolysaccharide [LPS] up-regulated genes and sites that increase with age: log_2_[OR] = 0.907, p = 7.03 x 10^-4^; overlap of LPS down-regulated genes and sites that decrease with age: log_2_[OR] = 1.715, p = 1.55 x 10^-3^)^27^. Our results thus suggest that the Amboseli baboon epigenetic clock not only tracks chronological age, but also captures age-related changes in DNA methylation levels that are functionally important for gene regulation.

### Comparison with other age-associated traits and differences between sexes

Overall, the clock performed favorably relative to other morphological or biomarker predictors of age in this population. The epigenetic clock generally explained more variance in true chronological age, resulted in lower median error, and exhibited less bias than predictions based on raw body mass index (BMI) or blood cell composition data from flow cytometry or blood smears (traits that change with age in baboons^28,29^). Its performance was comparable to molar dentine exposure, a classical marker of age^30^ (Supplementary Figure 6). For a subset of 30 individuals, we had two samples collected at different points in time. The predicted ages from these longitudinally collected samples were older for the later-collected samples, as expected (Figure 1C-D; binomial test p = 5.95 x 10^-5^). Furthermore, the change in epigenetic clock predictions between successive longitudinal samples positively predicted the actual change in age between sample dates (β = 0.312, p = 0.027, controlling for sex; difference between actual change and predicted change: mean 3.11 years ± 3.25 s.d.).

However, clock performance was not equivalent in males and females. Specifically, we observed that the clock was significantly more accurate in males (Figure 1; males: N = 135; MAD = 0.85 years ± 1.0 s.d.; Pearson’s *r* = 0.86, p = 5.49 x 10^-41^; females: N = 142; MAD = 1.6 years ± 2.4 s.d.; *r* = 0.78, p = 6.78 x 10^-30^; two-sided Wilcoxon test for differences in absolute error by sex: p = 4.37 x 10^-9^). Sex differences were also apparent in the slope of the relationship between predicted age and chronological age. Males show a 2.2-fold higher rate of change in predicted age, as a function of chronological age, compared to females (Figure 1A-B; chronological age by sex interaction in a linear model for predicted age: β = 0.448, p = 9.66 x 10^-19^, N = 277). Interestingly, sex differences are not apparent in animals < 8 years, which roughly corresponds to the age at which the majority of males have achieved adult dominance rank and dispersed from their natal group^31–33^ (N = 158, chronological age by sex interaction β = −0.038, p = 0.808). Rather, sex differences become apparent after baboons have reached full physiological and social adulthood (N = 119, chronological age by sex interaction β = 0.459, p = 9.74 x 10^-7^ in animals 8 years), when divergence between male and female life history strategies is most marked^31–33^ and when aging rates between the sexes are predicted to diverge^34–36^.

Because of these differences, we separated males and females for all subsequent analyses. However, we note that the effects of age on DNA methylation levels at individual clock sites are highly correlated between the sexes (Pearson’s *r* = 0.91, p = 3.35 x 10^-204^), with effect sizes that are, on average, more precisely estimated in males (paired t-test p = 4.53 x 10^-74^ for standard errors of *β_age_*; Supplementary Figure 4). In other words, the sex differences in clock performance reflect changes in methylation that occur at the same CpG sites, but with higher variance in females. Lower accuracy in females compared to males therefore appears to result from the greater variability in DNA methylation change in older females (Figure 1).

### Socioenvironmental predictors of variance in biological aging

Although the baboon epigenetic clock is a good predictor of age overall, individuals were often predicted to be somewhat older or younger than their known chronological age. In humans and some model systems, the sign and magnitude of this deviation captures information about physiological decline and/or mortality risk beyond that contained in chronological age alone^15–17,37^.

To test whether this observation extends to wild baboons, we focused on four factors of known importance to fitness in the Amboseli population. First, we considered cumulative early adversity, which is a strong predictor of shortened lifespan and offspring survival for female baboons^38,39^. We measured cumulative adversity as a count of major adverse experiences suffered in early life, including low maternal social status, early life drought, a competing younger sibling, maternal loss, and high experienced population density (i.e., social group size). Second, we considered social bond strength in adulthood, which positively predicts longer adult lifespan in baboons, humans, and other wild social mammals^40–43^. Third, we considered dominance rank, which is a major determinant of access to mates, social partners, and other resources in baboons^40,44–46^. Finally, we considered body mass index (BMI), a measure of body condition that, in the Amboseli baboons, primarily reflects lean muscle mass (mean body fat percentages have been estimated at <2% in adult females and <9% in adult males)^47^. Because raw BMI (i.e., BMI not correcting for age) also tracks growth and development (increasing as baboons reach their prime and then declining thereafter^28^, Supplementary Figure 7; Pearson’s *r* in males between rank and raw BMI = −0.56, p = 6.38 x 10^-9^), we calculated BMI relative to the expected value for each animal’s age using piecewise regression, which also eliminates correlations between BMI and male rank (Pearson’s *r* = −0.070, p = 0.504). We refer to this adjusted measure of BMI as age-adjusted BMI.

Because high cumulative early adversity and low social bond strength are associated with increased mortality risk in the Amboseli baboons, we predicted that they would also be linked to increased epigenetic age. For rank and age-adjusted BMI, our predictions were less clear: improved resource access could conceivably slow biological aging, but increased investment in growth and reproduction (either through higher fertility in females or physical competition for rank in males) could also be energetically costly. To investigate these possibilities, we modeled the deviation between predicted age and known chronological age (Δ_age_) as a function of cumulative early adversity, ordinal dominance rank, age-adjusted BMI, and for females, social bond strength to other females. Social bond strength was not included in the model for males, as this measure was not available for a large proportion of males in this data set (53.8%). We also included chronological age as a predictor in the model, as epigenetic age tends to be systematically overpredicted for young individuals and underpredicted for old individuals (Figure 1A-B; this bias has been observed in both foundational work on epigenetic clocks^5^ and recent epigenetic clocks calibrated for rhesus macaques^48^, as well as for elastic net regression analyses more generally^49^). Including chronological age in the model, as previous studies have done^5,7^, controls for this compression effect. None of the predictor variables were strongly linearly correlated (all Pearson’s *r* < 0.35; Supplementary Table 4).

Surprisingly, despite being two of the strongest known predictors of lifespan in wild baboons, neither cumulative early life adversity nor social bond strength explain variation in Δ_age_ (Table 1). In contrast, high male dominance rank is strongly and significantly associated with larger values of Δ_age_ (β = −0.078, p = 7.39 x 10^-4^; Figure 2; Table 1; Supplementary Figure 8). Alpha males are predicted to be an average of 10.95 months older than their true chronological age—a difference that translates to 11.5% of a male baboon’s expected adult lifespan in Amboseli. In contrast, dominance rank did not predict Δ_age_ in females (p = 0.228; Table 1). Finally, age-adjusted BMI also predicted Δ_age_ in males (p = 6.33 x 10^-3^) but not in females (p = 0.682; Table 1). Despite the tendency for high-ranking males to have higher raw BMI due to increased muscle mass, the effects of rank and age-adjusted BMI in males are distinct. Specifically, modeling dominance rank after adjusting for raw BMI also produces a significant effect of rank on Δ_age_ in the same direction (p = 9.93 x 10^-4^; Supplementary Table 5), as does substituting the age-adjusted BMI measure for either raw BMI or the residuals of raw BMI after adjusting for dominance rank (rank p = 1.52 x 10^-2^ and p = 1.88 x 10^-4^ respectively; Supplementary Table 5). In contrast, BMI is only a significant predictor of male Δ_age_ when corrected for age (i.e., age-adjusted) and when rank is included in the same model (Table 1; Supplementary Table 5). Further, we obtain the same qualitative results if all low BMI males are removed from the sample (BMI < 41; this cut-off was used because it drops all young males who have clearly not reached full adult size; p = 7.14 x 10^-3^; Supplementary Table 5). Dropping these males also eliminates the age-raw BMI correlation (Pearson’s *r* = −0.16, p = 0.21).

**Table 1.** Predictors of Δ_age_^1^

| Covariate | $\beta$<br>(Female) | P-value<br>(Female) | $\beta$<br>(Male) | P-value<br>(Male) |
| --- | --- | --- | --- | --- |
| Intercept | **5.400** | **$1.33 \times 10^{-15}$** | **3.294** | **$1.19 \times 10^{-8}$** |
| Cumulative early adversity | -0.050 | 0.807 | -0.005 | 0.973 |
| Social bond strength | 0.382 | 0.164 | — | — |
| Dominance rank | 0.025 | 0.228 | **-0.078** | **$7.39 \times 10^{-4}$** |
| Age-adjusted BMI | 0.026 | 0.682 | **0.111** | **$6.33 \times 10^{-3}$** |
| Chronological age | **-0.699** | **$1.62 \times 10^{-28}$** | **-0.277** | **$8.36 \times 10^{-8}$** |

<sup>1</sup>Separate linear models for $\Delta_{\text{age}}$ were fit for females ( $N = 66$ ) and for males ( $N = 93$ ) for whom no data values were missing; social bond strength was not included in the model for males. Significant results are shown in bold.

**Figure 2.**
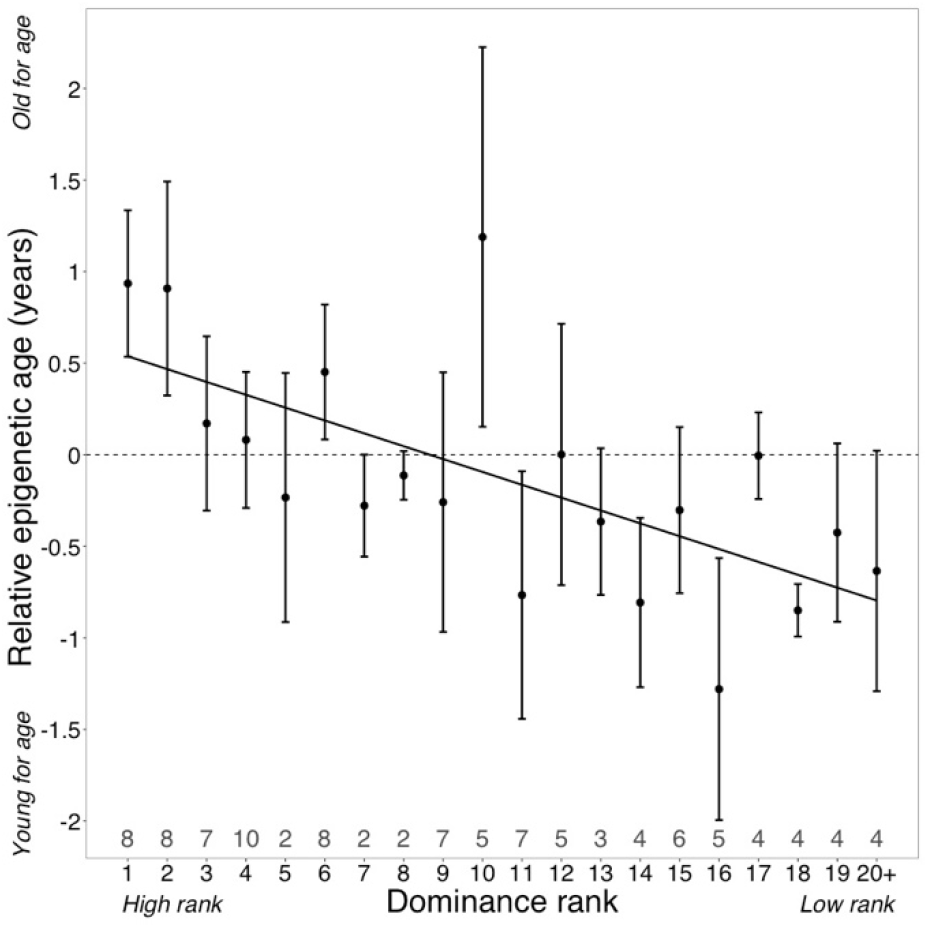
Dominance rank predicts relative epigenetic age in male baboons. High rank is associated with elevated values of Δ_age_ (β = −0.0785, p = 7.39 x 10^-4^, N = 105). The y-axis shows relative epigenetic age, a measure of epigenetic aging similar to Δ_age_ that is based on the sample-specific residuals from the relationship between predicted age and true chronological age. Positive (negative) values correspond to predicted ages that are older (younger) than expected for that chronological age. Dominance rank is measured using ordinal values, such that smaller values indicate higher rank. Dots and error bars represent the means and standard errors, respectively. Gray values above the x-axis indicate sample sizes for each rank.

### Male dominance rank predicts epigenetic age

In baboon males, achieving high rank depends on physical condition and fighting ability^33^. Consequently, rank in males is dynamic across the life course: males tend to attain their highest rank between 7 and 12 years of age and fall in rank thereafter (Supplementary Figure 9). Thus, nearly all males in the top four rank positions in our data set were between 7 and 12 years of age at the time they were sampled (however, because not all 7 – 12 year-olds are high– ranking, low rank positions include males across the age range; Supplementary Table 1, Supplementary Figure 9). We therefore asked whether the association between high rank in males and accelerated epigenetic aging is a function of absolute rank values, regardless of age, or deviations from the *expected* mean rank given a male’s age (i.e., “rank-for-age”; Supplementary Figure 9). We found that including rank-for-age as an additional covariate in the Δ_age_ model recapitulates the significant effect of ordinal male rank (p = 0.045), but finds no effect of rank-for-age (p = 0.819; Supplementary Table 5). Our results therefore imply that males incur the costs of high rank primarily in early to mid-adulthood, and only if they succeed in attaining high rank.

If attainment of high rank is linked to changes in epigenetic age within individuals, this pattern should be reflected in longitudinal samples. Specifically, males who improved in rank between samples should look older for age in their second sample relative to their first, and vice-versa. To assess this possibility, we calculated “relative epigenetic age” (the residuals of the best fit line relating chronological age and predicted age) for 14 males for whom we had repeated samples over time, 13 of whom changed ranks across sample dates (N = 28 samples, 2 per male). Samples collected when males were higher status predicted higher values of relative epigenetic age compared to samples collected when they were lower status (Figure 3; paired t-test, t = −2.99, p = 0.011). For example, in the case of a male whom we first sampled at low status (ordinal rank = 18) and then after he had attained the alpha position (ordinal rank 1), the actual time that elapsed between samples was 0.79 years, but he exhibited an increase in *predicted* age of 2.6 years. Moreover, the two males that showed a decrease in predicted age, despite increasing in chronological age (Figure 1D), were among those that experienced the greatest drop in social status between samples. Thus, change in rank between samples for the same male predicts change in Δ_age_, controlling for chronological age (R^2^ = 0.37, p = 0.021). Consistent with our cross-sectional results, we found a suggestive relationship between change in Δ_age_ and BMI (R^2^ = 0.31, p = 0.08). Here, too, the effect of dominance rank does not seem to be driven by BMI: while the association between change in Δ_age_ and change in rank is no longer significant when modeling rank after adjusting for raw BMI, the correlation remains consistent (R^2^ = 0.20, p = 0.167). In contrast, raw BMI adjusted for rank explains almost none of the variance in change in Δ_age_ (R^2^ = 0.01, p = 0.779).

**Figure 3.**
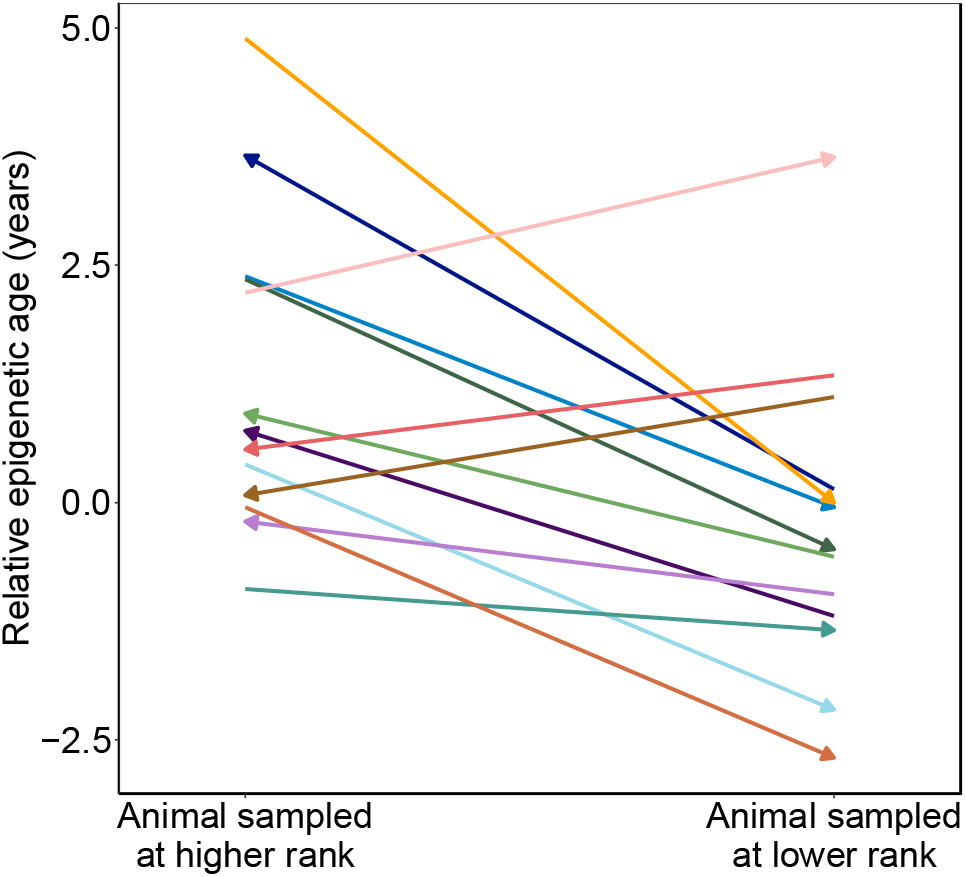
Male baboons exhibit higher relative epigenetic age when they occupy higher ranks. Relative epigenetic age for males in which multiple samples were collected when they occupied different ordinal rank values. Arrow indicates the temporal direction of rank changes: left-facing arrows represent cases in which the later sample was collected when males were higher-ranking, and right-facing arrows represent cases in which the later sample was collected when males were lower-ranking.

## Discussion

Together, our findings indicate that major environmental predictors of lifespan and mortality risk—particularly social bond strength and early life adversity in this population—do not necessarily predict epigenetic measures of biological age. Although this assumption is widespread in the literature, including for epigenetic clock analyses^50,51^, our results are broadly consistent with empirical results in humans. Specifically, while studies of early life adversity, which also predicts lifespan in human populations, find relatively consistent support for a relationship between early adversity and accelerated epigenetic aging in children and adolescents^12,13,52–56^, there is little evidence for the long-term effects of early adversity on epigenetic age in adulthood^14,57–61^. Thus, while DNA methylation may make an important contribution to the biological embedding of early adversity into adulthood^62,63^, it does not seem to do so through affecting the epigenetic clock itself. Social and environmental effects on the clock instead seem to be most influenced by concurrent conditions, lending support to “recency” models for environmental effects on aging that posit that health is more affected by the current environment than past experience^64–66^. Additional longitudinal sampling will be necessary to evaluate whether current conditions alone can explain accelerated epigenetic aging, or whether it also requires integrating the effects of exposures across the life course (the “accumulation” model^64,66^). Alternatively, the effects of early life adversity and social bond strength may act through entirely distinct pathways than those captured by our epigenetic clock (including targeting tissues or cell types that we were unable to assess here). Indeed, the proliferation of alternative epigenetic clocks in humans has revealed that the clocks that best predict chronological age are not necessarily the clocks that most closely track environmental exposures, and the same is likely to be true in other species^7,67^.

We found that the most robust socioenvironmental predictor of epigenetic age in the Amboseli baboons is male dominance rank, with a secondary effect of age-adjusted BMI observable when rank is included in the same model. Although high BMI also predicts accelerated epigenetic age in some human populations^37^, high BMI in these human populations is related to being overweight or obese. In contrast, because wild-feeding baboons in Amboseli are extremely lean^47^, the range of BMI in most human populations is distinct from the range exhibited by our study subjects (importantly, BMI in humans is calculated differently than BMI in baboons [see Methods], and therefore the BMI scales are species-specific). Instead, the higher BMI values in our dataset represent baboons in better body condition (more muscle mass). Given that rank in male baboons is determined by physical fighting ability^33^, these results suggest that investment in body condition incurs physiological costs that accelerate biological age. If so, the rank effect we observe may be better interpreted as a marker of competitiveness, not as a consequence of being in a “high rank” environment. In support of this idea, work on dominance rank and gene expression levels in the Amboseli baboons suggests that gene expression differences associated with male dominance rank tend to precede attainment of high rank, rather than being a consequence of behaviors exhibited after high rank is achieved^27^. Consistent with potential costs of attaining or maintaining high status, alpha males in Amboseli also exhibit elevated glucocorticoid levels^68^, increased expression of genes involved in innate immunity and inflammation^27^, and a trend towards elevated mortality risk^41^. Males who can tolerate these costs and maintain high rank are nevertheless likely to enjoy higher lifetime reproductive success, since high rank is the single best predictor of mating and paternity success in baboon males^33^.

This interpretation may also explain major sex differences in the effects of rank on epigenetic age, where dominance rank shows no detectable effect in females. Dominance rank in female baboons is determined by nepotism, not physical competition: females typically insert into rank hierarchies directly below their mothers, and hierarchies therefore tend to remain stable over time (and even intergenerationally)^69^. Our results contribute to an emerging picture in which dominance rank effects on both physiological and demographic outcomes are asymmetrical across sexes, and larger in males. Specifically, in addition to Δ_age_, male rank is a better predictor of immune cell gene expression and glucocorticoid levels than female rank^27,68,70^. Recent findings suggest that high rank may also predict increased mortality risk in male Amboseli baboons, whereas neither high nor low rank predicts increased mortality risk in females^41^. Together, these results argue that social status/dominance rank effects should not be interpreted as a universal phenomenon. Instead, the manner through which social status is achieved and maintained is likely to be key to understanding its consequences for physiology, health, and fitness^71^. Specifically, we predict that high status will be most likely to accelerate the aging process, including epigenetic age, in species-sex combinations where high status increases reproductive success or fecundity, and achieving status is energetically costly (e.g., male red deer, mandrills, and geladas; female meerkats^72–74^). Expanding studies of biological aging to a broader set of natural populations, especially those for which behavioral and demographic data are also available, will be key to testing these predictions.

## Methods

### Study population and biological sample collection

This study focused on a longitudinally monitored population of wild baboons *(Papio cynocephalus,* the yellow baboon, with some admixture from the closely related anubis baboon *P. anubis^75,76^*) in the Amboseli ecosystem of Kenya. This population has been continuously monitored by the Amboseli Baboon Research Project (ABRP) since 1971^22^. For the majority of study subjects (N = 242 of 245 individuals), birth dates were therefore known to within a few days’ error; for the remaining 3 individuals, birth dates were known within 3 months’ error (Supplementary Table 1).

All DNA methylation data were generated from blood-derived DNA obtained during periodic darting efforts, as detailed in^27,77,78^. Samples were obtained under approval from the Institutional Animal Care and Use Committee (IACUC) of Duke University and adhered to all the laws and regulations of Kenya. In brief, individually recognized study subjects were temporarily anesthetized using a Telazol-loaded dart delivered through a blow gun. Baboons were then safely moved to a new location where blood samples and morphometric data, including body mass and crown-rump length, were collected. Baboons were then allowed to recover from anesthesia in a covered holding cage and released to their group within 2 – 4 hours. Blood samples were stored at −20° C in Kenya until export to the United States.

### DNA methylation data

DNA methylation data were generated from blood-extracted DNA collected from known individuals in the Amboseli study population (N = 277 samples from 245 animals; 13 females and 15 males were each sampled twice, and 1 female and 1 male were each sampled three times). Here, we analyzed a combined data set that included previously published reduced representation bisulfite sequencing^79^ (RRBS) data from the same population (N = 36 samples)^23^ and new RRBS data from 241 additional samples.

RRBS libraries were constructed following^80^, using ~200 ng baboon DNA plus 0.2 ng unmethylated lambda phage DNA per sample as input. Samples were sequenced to a mean depth of 17.8 (± 10.5 s.d.) million reads on either the Illumina HiSeq 2000 or HiSeq 4000 platform (Supplementary Table 1), with an estimated mean bisulfite conversion efficiency (based on the conversion rate of lambda phage DNA) of 99.8% (minimum = 98.1%). Sequence reads were trimmed with Trim Galore! to remove adapters and low quality sequence (Phred score < 20). Trimmed reads were mapped with BSMAP^82^ to the baboon genome *(Panu2.0)* allowing a 10% mismatch rate to account for the degenerate composition of bisulfite-converted DNA. We used the mapped reads to count the number of methylated and total reads per CpG site, per sample^82^. Following ^23,25^, CpG sites were filtered to retain sites with a mean methylation level between 0.1 and 0.9 (i.e., to exclude constitutively hyper- or hypo-methylated sites) and mean coverage ≥5x. We also excluded any CpG sites with missing data for ≥ 5% of individuals in the sample. After filtering, we retained N = 458,504 CpG sites for downstream analysis. For the remaining missing data (mean number of missing sites per sample = 1.4% ± 3.5% s.d., equivalent to 6,409 ± 16,024 s.d. sites), we imputed methylation levels using a k-nearest neighbors approach in the R package *impute,* using default parameters^83^.

### Building the epigenetic clock

We used the R package *glmnet*^84^ version 2.0.10 to build a DNA methylation clock for baboons. Specifically, we fit a linear model in which the predictor variables were normalized levels of DNA methylation at 458,504 candidate clock CpG sites across the genome and the response variable was chronological age. To account for the excess of CpG sites relative to samples, *glmnet* uses an elastic net penalty to shrink predictor coefficients toward 0^85^. Optimal alpha parameters were identified by grid searching across a range of alphas from 0 (equivalent to ridge regression) to 1 (equivalent to Lasso) by increments of 0.1, which impacts the number of clock CpG sites by varying the degree of shrinkage of the predictor coefficients toward 0 (Supplementary Figure 2). We defined the optimal alpha as the value that maximized R between predicted and true chronological age across all samples. We set the regularization parameter lambda to the value that minimized mean-squared error during n-fold internal cross-validation.

To generate predicted age estimates for a given sample, we used a leave-one-out cross-validation approach in which all samples but the “test” sample were included for model training, and the resulting model was used to predict age for the left-out test sample. Importantly, training samples were scaled independently of the test sample in each leave-one-out model to avoid bleed-through of information from the test data into the training data. To do so, we first quantile normalized methylation ratios (the proportion of methylated counts to total counts for each CpG site) within each sample to a standard normal distribution. Training samples were then separated from the test sample and the methylation levels for each CpG site in the training set were quantile normalized across samples to a standard normal distribution. For predicting age in the test sample, we compared the methylation value for each site in the test sample to the empirical cumulative distribution function for the training samples (at the same site) to estimate the quantile in which the training sample methylation ratio fell. The training sample was then assigned the same quantile value from the standard normal distribution using the function *qnorm* in R.

### Epigenetic clock enrichment analyses

To evaluate whether CpG sites included in the epigenetic clock were enriched in functionally important regions of the baboon genome^25,86^, we used two-sided Fisher’s exact tests to investigate enrichment/depletion of the 573 epigenetic clock sites in (i) gene bodies and exons, based on the Ensembl annotation *Panu2.0.90;* (ii) CpG islands annotated in the UCSC Genome Browser; (iii) CpG shores, defined as the 2,000 basepairs flanking CpG islands (following^25,86,87^); and (iv) promoter regions, defined as the 2,000 basepairs upstream of the 5’-most annotated transcription start site for each gene (following ^25,86^). We also considered (v) putative enhancer regions, which have not been annotated for the *Panu2.0* assembly. We therefore used ENCODE H3K4me1 ChIP-seq data from humans^88^ and the *liftOver* tool to define likely enhancer coordinates in *Panu2.0.*

We also tested for enrichment of clock sites in regions of the genome that have been identified by previous empirical studies to be of special interest. First, we considered regions that likely have regulatory activity in blood cells, defined as all 200 base-pair windows that showed evidence of enhancer activity in a recently performed massively parallel reporter assay^26^. We used *liftOver* to identify the inferred homologous *Panu2.0* coordinates for these windows, which were originally defined in the human genome. Second, we defined age-related differentially methylated regions (age DMRs) in the Amboseli baboons based on genomic intervals found, in previous analyses, to contain at least three closely spaced age-associated CpG sites (inter-CpG distance ≤ 1kb), as described in ^25^. Third, because inflammatory processes involved in innate immunity are strongly implicated in the aging process, we defined lipopolysaccharide (LPS) up- regulated and LPS down-regulated genes as those genes that were significantly differentially expressed (1% false discovery rate) between unstimulated Amboseli baboon white blood cells and LPS-stimulated cells from the same individual, following 10 hours of culture in parallel^27^.

### Comparisons to alternative predictors of aging

To assess the utility of the DNA methylation clock relative to other data types, we compared its predictive accuracy to clocks based on three other age-related phenotypes: tooth wear (percent molar dentine exposure^30^), body condition (body mass index: BMI^28^), and blood cell type composition (blood smear counts and lymphocyte/monocyte proportions from flow cytometry performed on peripheral blood mononuclear cells, as in ^27,89^). Leave-one-out model training and prediction were performed for each data type using linear modeling. To compare the relative predictive accuracy of each data type, we calculated the R^2^ between predicted and chronological age, the median absolute difference between predicted and chronological age, and the bias in age predictions (the absolute value of 1 – slope of the best fit line between predicted and chronological age) (Supplementary Figure 6).

#### Tooth wear

Molar enamel in baboons wears away with age to expose the underlying dentine layer. Percent dentine exposure (PDE) on the molar occlusal surface has been shown to be strongly age-correlated in previous work^30^. To assess its predictive power, we obtained PDE data from tooth casts reported by Galbany and colleagues^30^ for the left upper molars (tooth positions M1, M2, M3) and left lower molars (tooth positions M1, M2, M3) for 39 males and 34 females in our data set. For each molar position (M1, M2, M3) within each individual, we calculated PDE as the mean for the upper and lower molars. Because dentine exposure scales quadratically with respect to age, we fit age as a function of PDE using the following model:

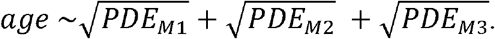

#### Body mass index

For both male and female baboons in Amboseli, body mass increases with age until individuals reach peak size, and then tends to decrease with age as animals lose fat and/or muscle mass^28^. To quantify body condition using body mass, we calculated body mass index (BMI) values for 139 males and 154 females for whom body mass and crown-rump length data were available from periodic darting efforts. We retained only measures taken from animals born into and sampled in wild-feeding study groups, when sex-skin swellings (in females only) that could affect crown-rump length measures were absent. BMI was calculated as mass (kilograms) divided by crown-rump length (meters squared), following^47^. To assess the predictive power of age-adjusted BMI, we built sex-specific piecewise-regression models using the package *segmented* in R^90^. Breakpoints for the piecewise-regression models (to separate “youthful” versus “aged” animals) were initialized at 8 years old for males and 10 years old for females, following findings from previous work on body mass in the Amboseli population^28^.

#### Blood cell type composition

The proportions of different cell types in blood change across the life course, including in baboons^29^. We assessed the predictive power of blood cell composition for age using two data sets. First, we used data collected from blood smear counts (N = 134) for five major white blood cell types: basophils, eosinophils, monocytes, lymphocytes, and neutrophils. Second, we used data on the proportional representation of five peripheral blood mononuclear cell (PBMC) subsets: cytotoxic T cells, helper T cells, B cells, monocytes, and natural killer cells, measured using flow cytometry as reported by Lea and colleagues^27^ (N = 53). Cell types were included as individual covariates for leave-one-out model training.

### Sources of variance in predicted age

We asked whether factors known to be associated with inter-individual variation in fertility or survival also predict inter-individual variation in Δ_age_ (predicted age from the epigenetic clock minus known chronological age). To do so, we fit linear models separately for males and females, with Δ_age_ as the dependent variable and dominance rank at the time of sampling, cumulative early adversity, age-adjusted BMI, and chronological age as predictor variables^38^. For females, we also included a measure of social bond strength to other females as a predictor variable, based on findings that show that socially isolated females experience higher mortality rates in adulthood^40,91^. Samples with missing values for any of the predictor variables were excluded in the model, resulting in a final analysis set of 66 female samples (from 59 females) and 93 male samples (from 84 males). The chronological ages of samples with complete data relative to samples with missing data were equivalent for females (t-test, t = 1.95, p = 0.053) but were slightly lower for males (t-test, t = −3.04, p = 0.003; mean chronological ages are 7.98 and 9.65 years for complete and missing samples, respectively). Predictor variables were measured as follows.

#### Dominance rank

Sex-specific dominance hierarchies were constructed monthly for every social group in the study population based on the outcomes of dyadic agonistic encounters. An animal was considered to win a dyadic agonistic encounter if it gave aggressive or neutral, but not submissive, gestures, and the other animal gave submissive gestures only^92^. These wins and losses were entered into a sex-specific data matrix, such that animals were ordered to minimize the number of entries falling below the matrix diagonal (which would indicate that the lower ranked individual won a dyadic encounter). Ordinal dominance ranks were assigned on a monthly basis to every adult based on these matrices, such that low numbers represent high rank/social status and high numbers represent low rank/social status^33,69^. Although most analyses of data from the Amboseli baboons have used ordinal ranks as the primary measure of social status, in some analyses proportional rank (i.e., the proportion of same-sex members of an individual’s social group that he or she dominates) has proven to be a stronger predictor of other trait outcomes^93,94^. In this study, we chose to use ordinal ranks, but proportional and ordinal dominance rank were highly correlated in this particular data set (R^2^ = 0.70, p = 1.13 x 10^-58^). Using ordinal rank rather than proportional rank therefore did not qualitatively affect our analyses. Additionally, to investigate whether the patterns we observed are due to a consistent effect of rank across all ages, or instead an effect of being high or low rank relative to the expected (mean) value for a male’s age, we also calculated a “rank-for-age” value. Rank-for-age is defined as the residuals of a model with dominance rank as the response variable and age and age^2^ as the predictor variables (Supplementary Figure 8).

#### Cumulative early adversity

Previous work in Amboseli defined a cumulative early adversity score as the sum of 6 different adverse conditions that a baboon could experience during early life^38^. This index strongly predicts adult lifespan in female baboons, and a modified version of this index also predicts offspring survival^39^. To maximize the sample size available for analysis, we excluded maternal social connectedness, the source of adversity with the highest frequency of missing data, leaving us with a cumulative early adversity score generated from 5 different binary-coded adverse experiences. These experiences were: (i) early life drought (defined as ≤ 200 mm of rainfall in the first year of life), which is linked to reduced fertility in females^46,95^; (ii) having a low ranking mother (defined as falling within the lowest quartile of ranks for individuals in the data set), which predicts age at maturation^96–98^; (iii) having a close-inage younger sibling (< 1.5 years), which may redirect maternal investment to the sibling^99^, (iv) being born into a large social group, which may increase within-group competition for shared resources^46,98,100^, and (v) maternal death before the age of 4, which results in a loss of both social and nutritional resources^98,101^.

#### Body mass index

Age-adjusted BMI was modeled as the residuals from sex-specific piecewise regression models relating raw BMI to age. By taking this approach, we asked whether having relatively high BMI for one’s age and sex predicted higher (or lower) Δ_age_. To calculate rank-adjusted BMI values, we modeled raw BMI as a function of rank in a linear model and calculated the residuals from the model. To calculate dominance rank adjusted for raw BMI, we took the inverse approach. We note that BMI for baboons is not directly comparable to BMI for humans because baboon BMI is measured as body mass divided by the square of crown-rump length (because baboons are quadrupedal), whereas human BMI is calculated as body mass divided by the square of standing height.

#### Social bond strength

For this analysis, we measured female social bond strength to other females using the dyadic sociality index (DSI_F_)^41^. We did not include this parameter (male’s social bond strength to females) for the male model, because this measure is unavailable for many males in this data set. DSI_F_ was calculated as an individual’s average bond strength with her top three female social partners, in the 365 days prior to the day of sampling, controlling for observer effort. This approach is based on representative interaction sampling of grooming interactions between females, in which observers record all grooming interactions in their line of sight while moving through the group conducting random-ordered, 10-minute long focal animal samples of pre-selected individuals. Because smaller groups receive more observer effort per individual and per dyad (and thus record more grooming interactions per individual or dyad), we estimated observer effort for dyad *d* in year*y* as:

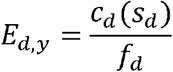

where *c_d_* is the number of days the two females in a dyad were coresident in the same social group, *s_d_* is the number of focal samples taken during the dyad’s coresidence, and *f_d_* is the average number of females in the group during the dyad’s coresidence.

DSI_F_ for each adult female dyad in each year is the z-scored residual, ε, from the model:

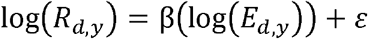

where *R_d,y_* is the number of grooming interactions for dyad *d* in year *y* divided by the number of days that the two individuals were coresident, and *E_d,y_* is observer effort.

### Analysis of longitudinal samples

To test whether changes in rank predict changes in relative epigenetic age within individuals, we used data from 11 males from the original data set and generated additional RRBS data for 9 samples, resulting in a final set of 14 males who each were sampled at least twice in the data set, 13 of whom occupied different ordinal ranks at different sampling dates (mean years elapsed between samples = 3.7 ± 1.65 s.d.; mean absolute difference in dominance ranks = 1.29 ± 8.34 s.d.). This effort increased our total sample size to N = 286 samples from 248 unique individuals. To incorporate the new samples into our analysis, we reperformed leave-one-out age prediction with N-fold internal cross validation at the optimal alpha selected for the original N = 277 samples (alpha = 0.1). For the 277 samples carried over from the original analysis, we verified that age predictions were nearly identical between the previous analysis and the expanded data set (R^2^ = 0.98, p = 2.21 x 10^-239^; Supplementary Table 1).

Based on the new age predictions for males in the data set (N = 140), we again calculated relative epigenetic age as the residual of the best fit line relating predicted age to chronological age. We then used the 14 males with repeated DNA methylation profiles and rank measures in this data set to test whether, within individuals, changes in dominance rank or rank-for-age explained changes in relative epigenetic age between samples. In total, five males were sampled three times. For four of these five, we only included the two samples that were sampled the farthest apart in time (i.e., excluded the temporal middle sample) to maximize the age change between sample dates. For the fifth male, BMI information was missing for the third sample, so we included the first two samples collected in time.

## Data Availability

All sequencing data generated during this study are available in the NCBI Sequence Read Archive (project accession PRJNA648767; reviewer access: ##########).

## Code Availability

All R code used to analyze data in this study are available at https://github.com/janderson94/BaboonEpigeneticAging.

## Acknowledgements

We gratefully acknowledge the support provided by the National Science Foundation and the National Institutes of Health for the majority of the data represented here, currently through NSF IOS 1456832, NIH R01AG053308, R01AG053330, R01HD088558, and P01AG031719. R.A.J. is supported by NIH F32HD095616 and J.A.A. by NSF #2018264636. We also acknowledge support for high-performance computing resources from the North Carolina Biotechnology Center (Grant Number 2016-IDG-1013) and a seed grant from the Center for Population Health and Aging (P30AG034424 to A. O’Rand). We thank the members of the Amboseli Baboon Research Project for collecting the data presented here, especially J. Altmann for her foundational role in establishing the study population and these data sets; J. Gordon, N. Learn, and K. Pinc for managing the database; R.S. Mututua, S. Sayialel, and J.K. Warutere for data collection in the field; and T. Wango and V. Oudu for their assistance in Nairobi. We also thank the Kenya Wildlife Service, University of Nairobi, the Institute of Primate Research, the National Museums of Kenya, the National Council for Science, Technology, and Innovation, members of the Amboseli-Longido pastoralist communities, the Enduimet Wildlife Management Area, Ker & Downey Safaris, Air Kenya, and Safarilink for their assistance in Kenya. Finally, we thank J. Galbany for assistance with the molar dentine data set; current and past members of the Tung, Alberts, Archie, and Altmann labs for their helpful feedback; and J. Higham and two anonymous reviewers for constructive critiques of a previous version of this manuscript. This research was approved by IACUCs at Duke University, University of Notre Dame, and Princeton University and adhered to all the laws and regulations of Kenya. For a complete set of acknowledgments of funding sources, logistical assistance, and data collection and management, please visit http://amboselibaboons.nd.edu/acknowledgements/.

## Author Contributions

Conceptualization, R.A.J., J.A.A., J.T., E.A.A., A.J.L.; Investigation, J.A.A., R.A.J., A.J.L., F.A.C., M.Y.A., T.N.V., and J.T.; Formal Analysis, J.A.A. and R.A.J.; Writing—Original Draft, R.A.J., J.A.A., and J.T.; Writing—Reviewing & Editing, R.A.J., J.A.A., A.J.L., T.N.V., M.Y.A., F.A.C., S.C.A., E.A.A., and J.T. Funding Acquisition, J.T., S.C.A., and E.A.A. Supervision, J.T.

---

**Supplementary Table 4.** Pearson correlations among covariates for females (above diagonal) and males (below diagonal), with p-values in parentheses.

|  | cumulative | DSI<sub>F</sub> | rank | Age-adjusted BMI | age |
| --- | --- | --- | --- | --- | --- |
| cumulative | - | -0.222 (0.073) | 0.310 (0.011) | 0.098 (0.432) | -0.284 (0.021) |
| DSI<sub>F</sub> | NA | - | -0.266 (0.031) | -0.188 (0.131) | 0.112 (0.372) |
| rank | -0.058 (0.578) | NA | - | 0.058 (0.646) | 0.218 (0.078) |
| Age-adjusted BMI | -0.038 (0.719) | NA | -0.068 (0.516) | - | -0.098 (0.434) |
| age | 0.133 (0.202) | NA | -0.313 (0.002) | -0.075 (0.476) | - |

**Supplementary Figure 1.**
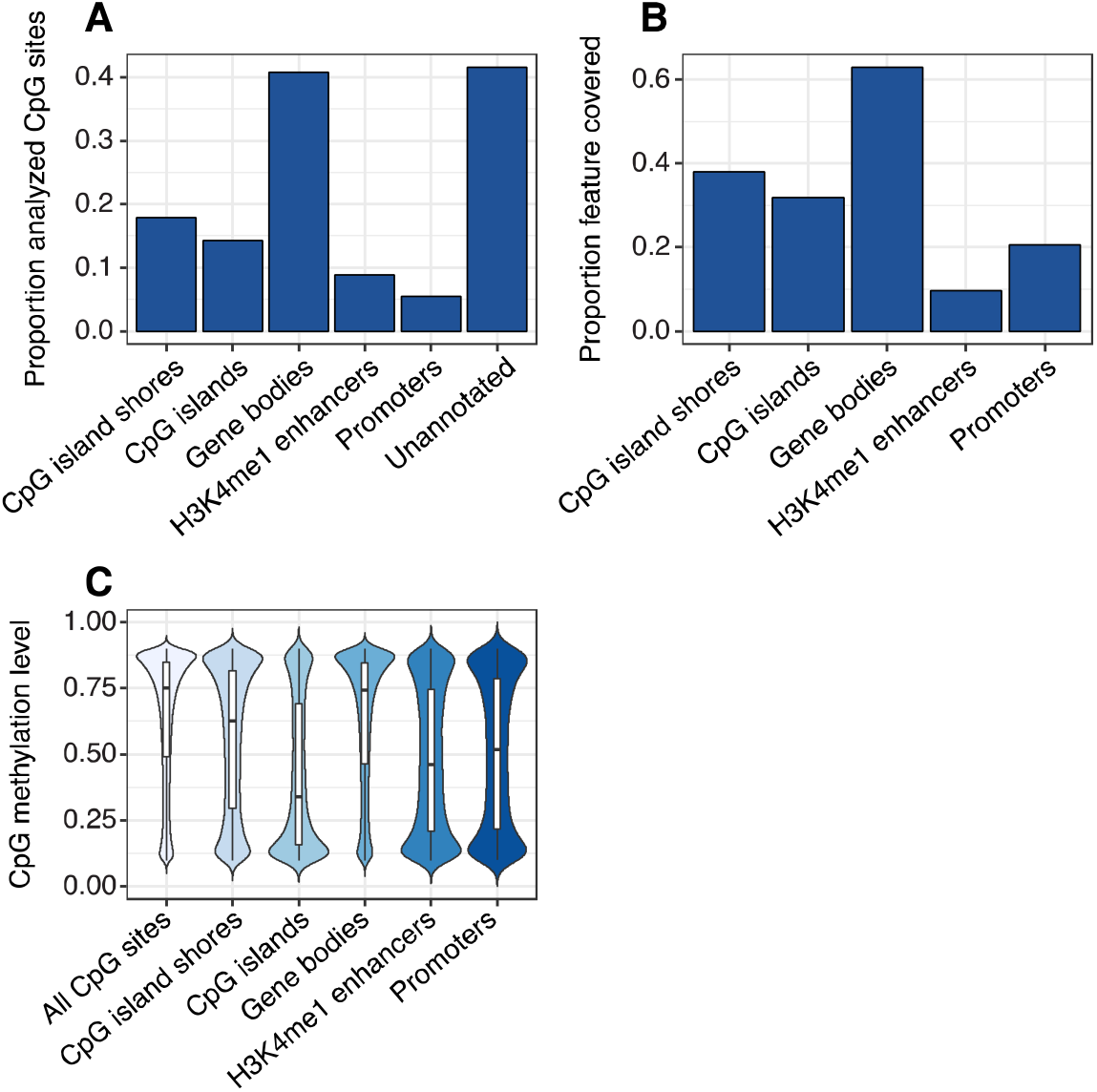
Characteristics of the RRBS data set. **(A)** Proportion of the 458,504 evaluated CpG sites that overlapped annotated features of the *Panu2* genome. **(B)** Proportion of annotated features in the *Panu2* genome that overlapped at least one of the 458,504 evaluated CpG sites. **(C)** Distribution of mean DNA methylation levels for CpG sites within annotated features of the *Panu2* genome. Each white box represents the interquartile range, with the median value depicted as a black horizontal bar. Whiskers extend to the most extreme values within 1.5 x the interquartile range. As expected, CpG sites tended to be highly methylated genome-wide and have lower average methylation in promoters, enhancers, and CpG islands.

**Supplementary Figure 2.**
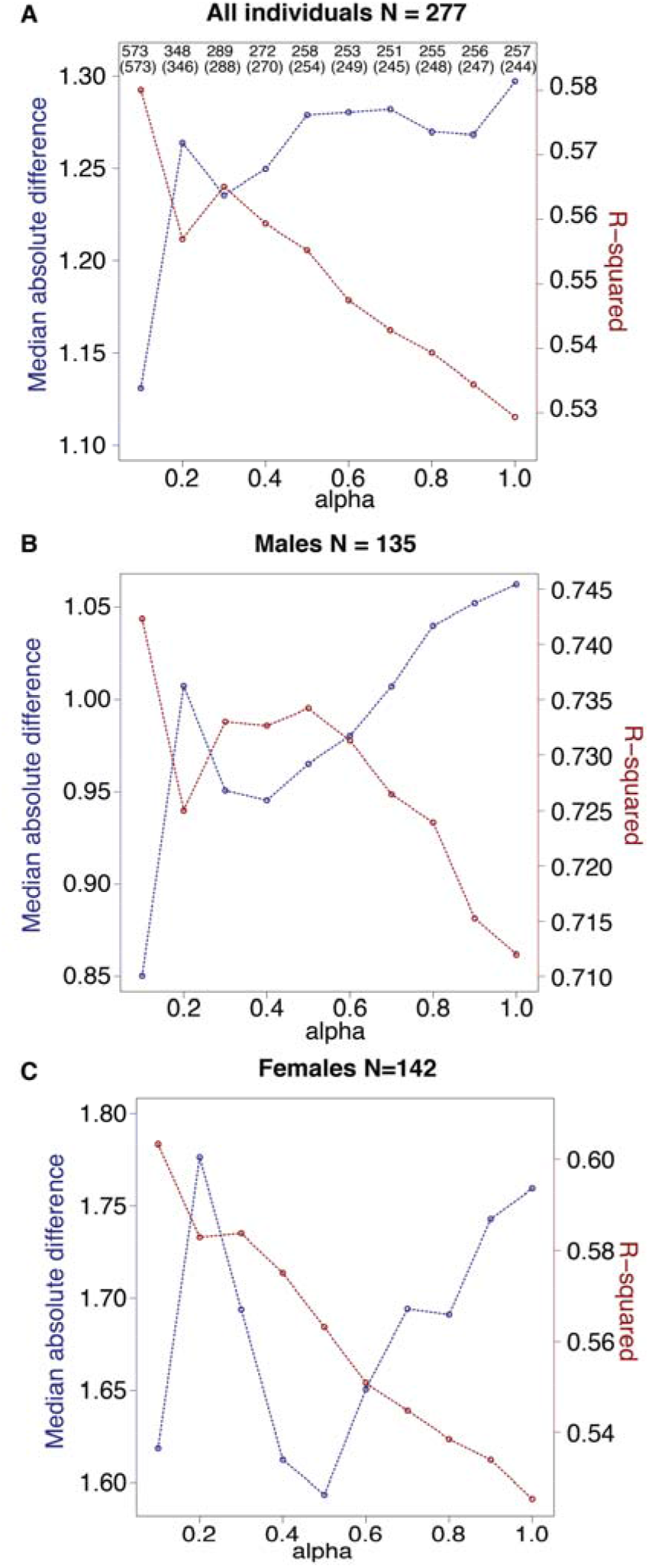
Comparison of clock performance across alternative values of alpha. Alpha was set via grid search across possible values from 0.1 to 1, in steps of 0.1, and chosen based on the highest R^2^ value between predicted age and known chronological age (red lines). The blue lines show the median absolute difference between predicted and true age (lower is better), and exhibits roughly inverse behavior to R^2^. **(A)** For each clock generated with a different alpha value, the total number of CpG sites included in the clock is shown on top, and the number of clock sites that overlap the final clock used in this study (N = 573 sites, alpha = 0.1) is given in parentheses immediately below. **(B, C)** As in (A), but with results shown specifically for males **(B)** versus females **(C)**.

**Supplementary Figure 3.**
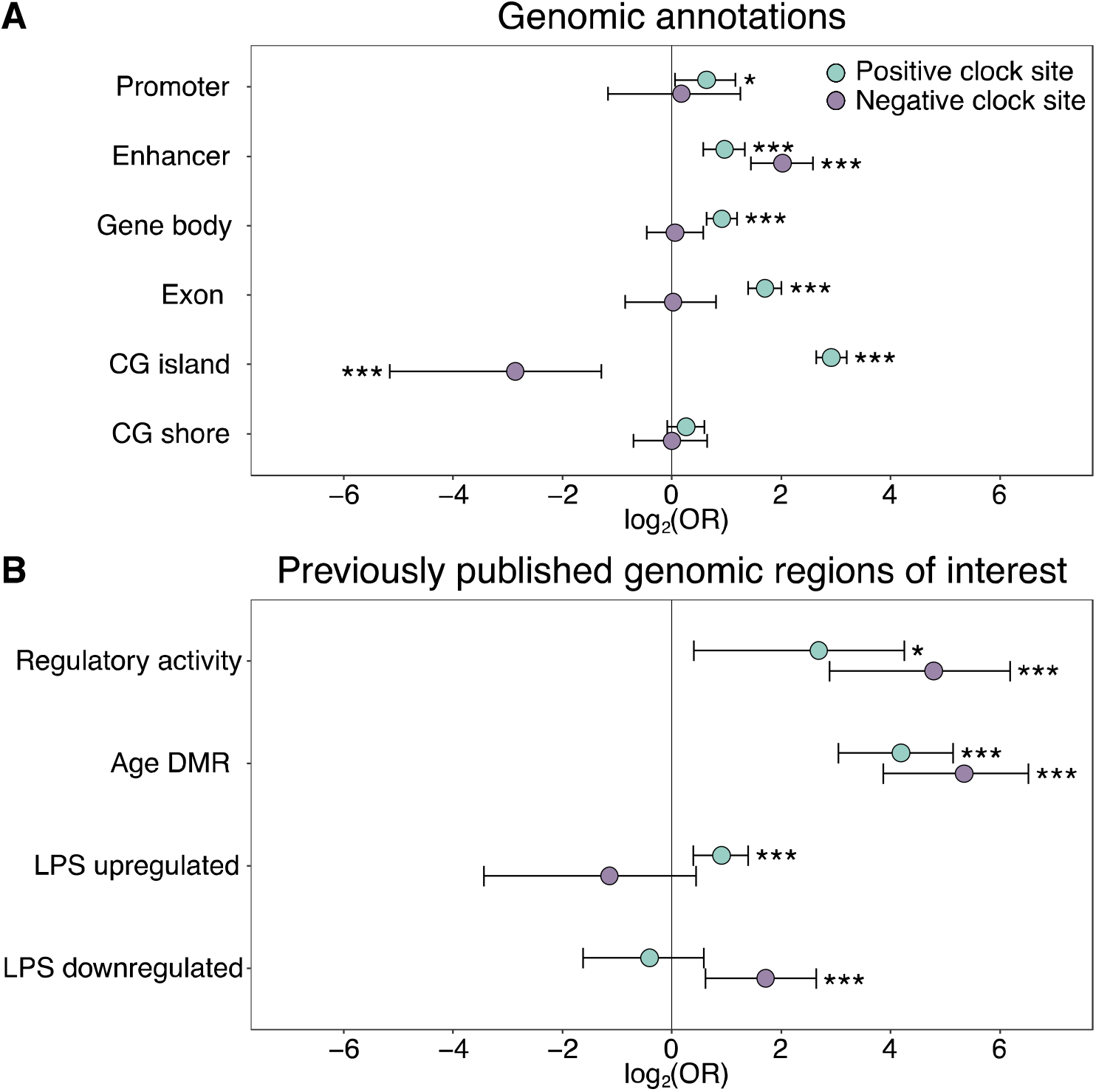
Enrichment of the epigenetic clock CpG sites by genomic compartment. The log_2_(odds ratio) of CpG sites in the epigenetic clock, relative to all 458,504 CpG sites initially evaluated, in **(A)** annotated genomic regions and **(B)** in loci with putative regulatory activity or in or near genes that are responsive to age or immune stimulation. Regions of regulatory activity were identified with the massively parallel reporter assay, mSTARR-Seq^26^, following a liftover from the human genome to the baboon genome to identify putatively orthologous coordinates. Age differentially methylated regions (DMR) and genes responsive to lipopolysaccharide (LPS) were previously identified from blood samples from the same baboon population^25,27^. Two-sided Fisher’s exact tests were performed separately for epigenetic clock sites that increased (positive clock sites: N = 459) or decreased (negative clock sites: N = 134) in DNA methylation levels with age. See Supplementary Table 2 for a complete list of the genomic locations of the 573 epigenetic clock sites. * p < 0.05, *** p < 0.005.

**Supplementary Figure 4.**
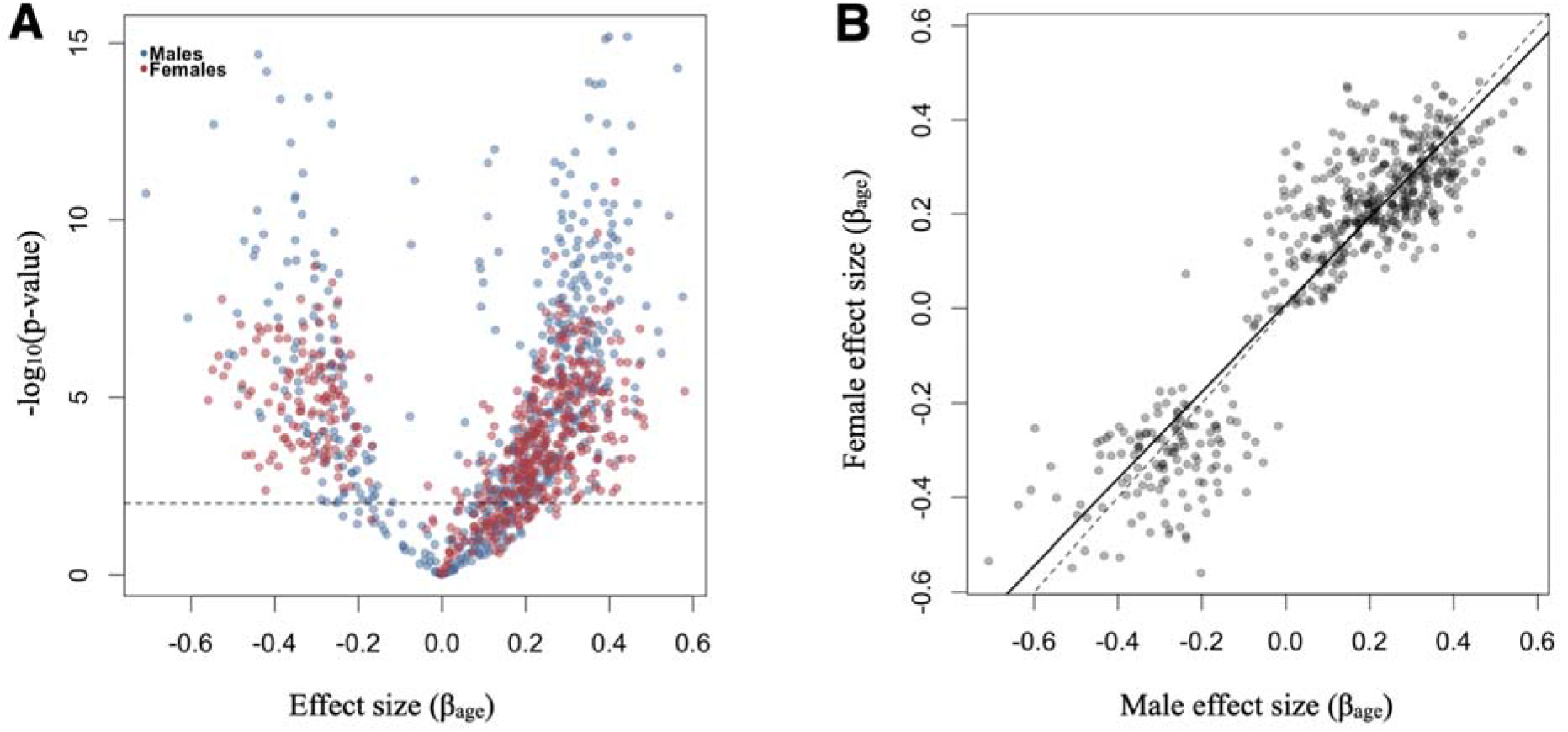
Association between age and DNA methylation level for individual clock CpG sites. **(A)** Volcano plot of the effect size (*β*_age_) versus the -log_10_(p-value) of age effects on DNA methylation for males (blue) and females (red), based on estimates from **a** binomial mixed-effects model designed for bisulfite sequencing data^25^. Results for the 534 sites that could be modeled using this approach are shown. Other predictor variables in the model included a fixed effect for sample batch and a random effect that controlled for kinship (estimated via Queller and Goodnight’s *r* and multilocus microsatellite genotype data in the program *coancestry^102^*). Dashed line corresponds to a nominal p-value of 0.01. **(B)** Age effects on DNA methylation estimated separately in males and females are highly correlated (R^2^ = 0.83, p = 3.35 x 10^-204^). The dashed line indicates the y = x line. The solid black line indicates the best fit line.

**Supplementary Figure 5.**
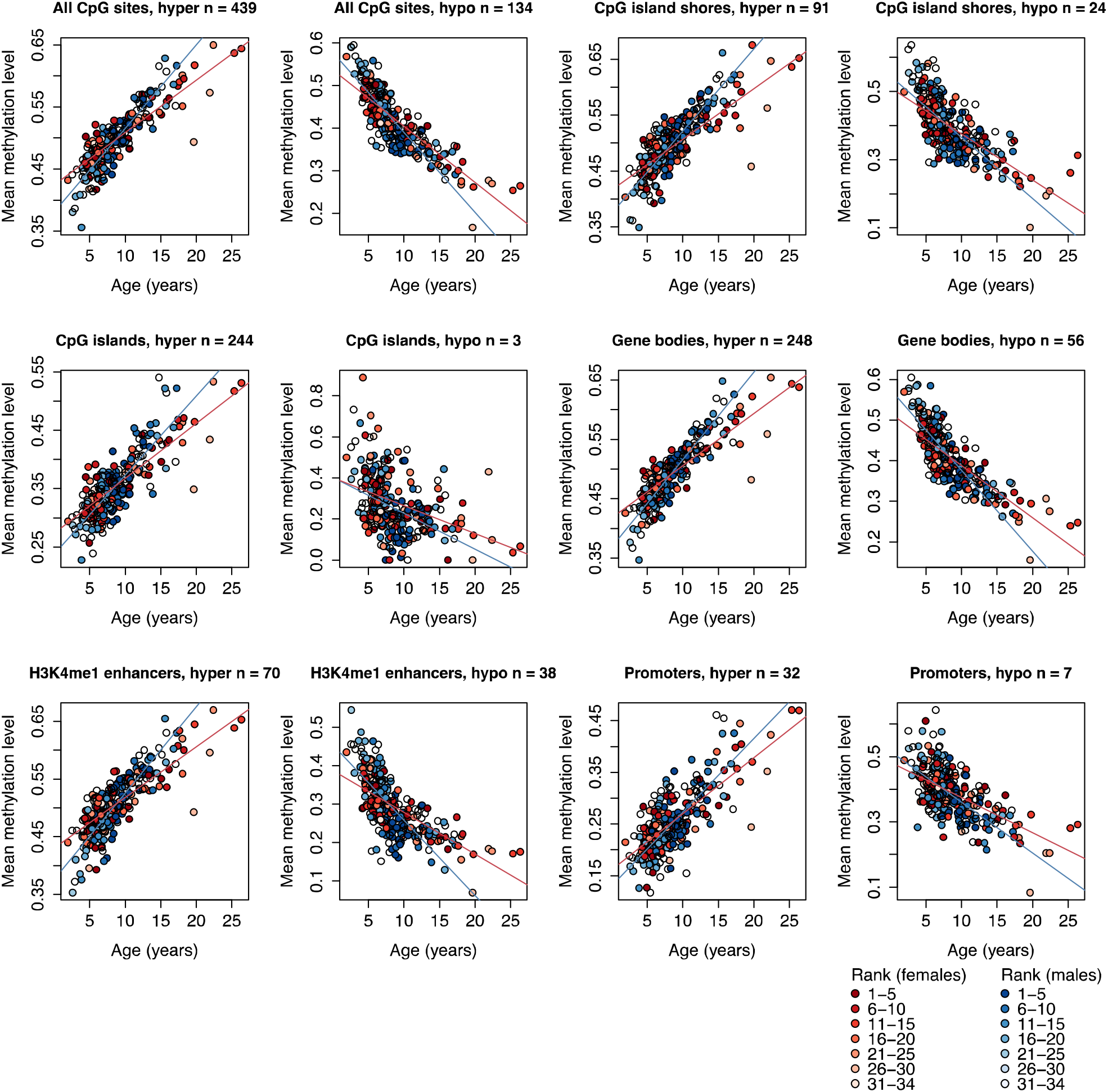
Methylation levels of clock CpG sites across different genomic compartments. Each circle represents a sample, with chronological age of the animal at time of sampling shown on the x-axis. The y-axis represents the average methylation level for that sample across CpG clock sites that overlap the annotated genomic region shown in the panel label, stratified by sites that increased (denoted “hyper”) or decreased (denoted “hypo”) methylation levels with age. Number of clock sites overlapping each annotated region is given in each panel title; a clock site can overlap multiple annotated regions, and can therefore be represented in more than one plot. Red and blue lines represent best fit lines for female and male samples, respectively. All best fit lines are significant (p < 1 x 10^-4^).

**Supplementary Figure 6.**
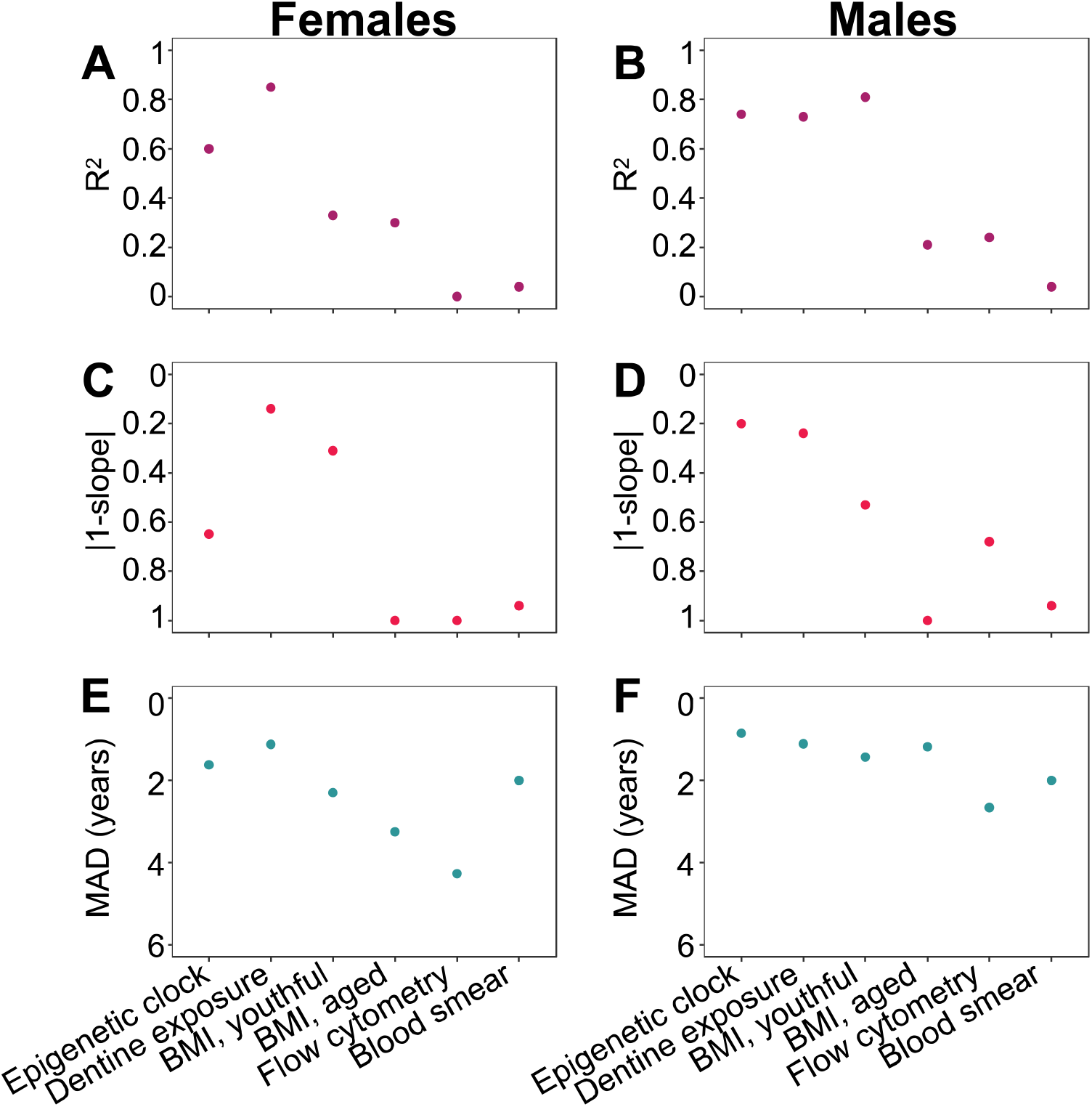
Comparison of the performance of the epigenetic clock to other predictors of chronological age. Performance measures of age predictors are presented separately for females **(A, C, E)** and males **(B, D, F)** except for differential white blood cell counts (blood smears), where males and females were combined. Predictors are ordered in the same fashion in all panels (epigenetic clock to the left, and then following highest to lowest R^2^ in females). The breakpoint to define youthful versus aged animal BMI was 10 and 8 years old for females and males, respectively. **(A-B)** Adjusted R^2^ between predicted age and true chronological age. **(C-D)** Absolute difference between the y = x line (slope of one) and the slope of the best-fit line of predicted age as a function of true chronological age. This metric captures bias in age prediction estimates (values that are lower on the reverse-coded y-axis are more biased). **(E-F)** Median absolute difference (MAD) between each individual’s predicted age and true chronological age (values that are lower on the reverse-coded y-axis have higher MAD).

**Supplementary Figure 7.**
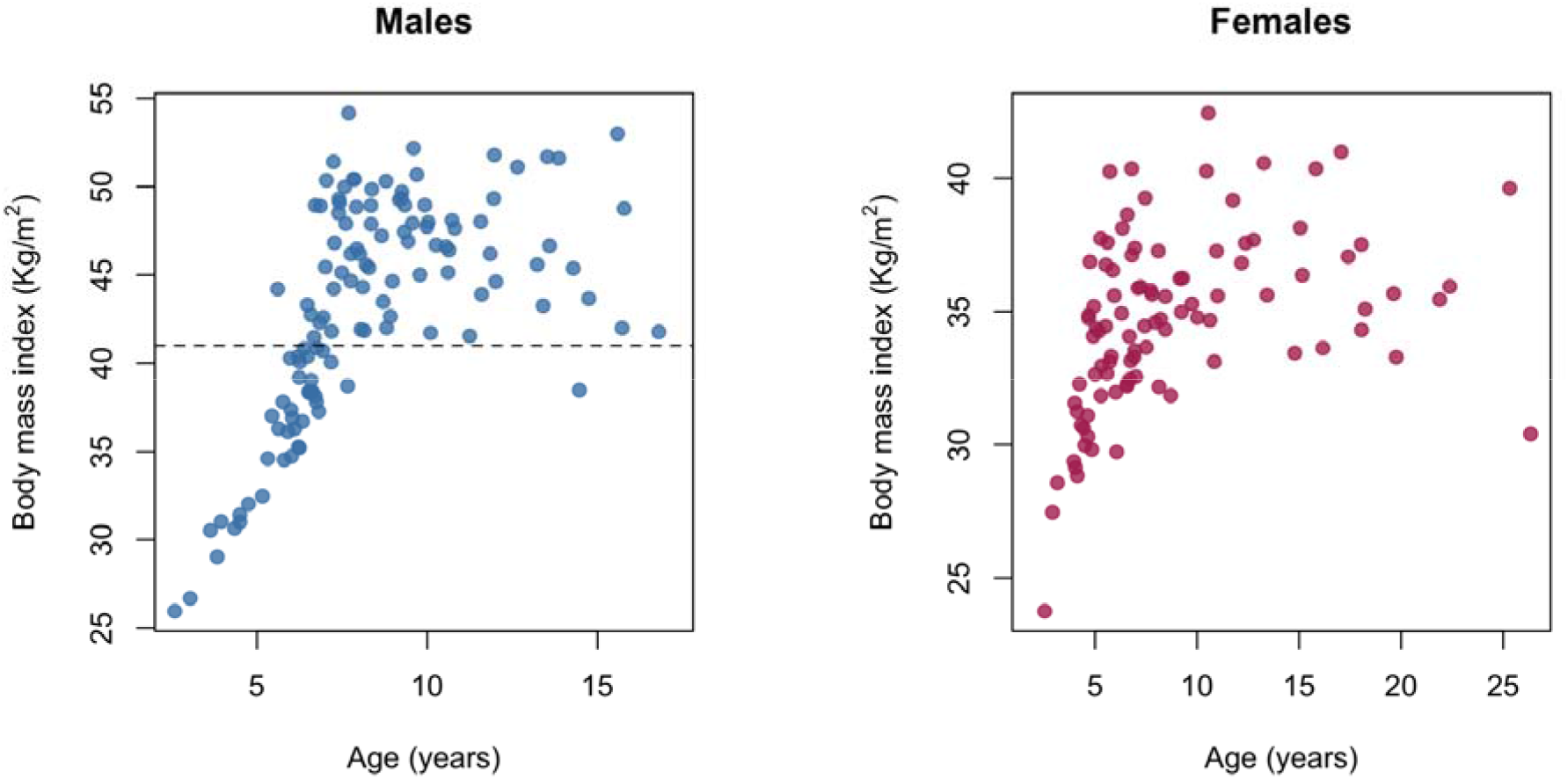
The relationship between age and body mass index in the Amboseli baboons. Chronological age in years at the time of sampling versus body mass index (kilograms/meters^2^) for males and females in our sample. Two distinct patterns are observable for both sexes: a stage when animals are still growing (prior to ~7 – 8 years old) and a stage in which animals vary in BMI as adults. BMI in baboons is measured using the distance between the crown of the head and the rump as the “height” measure, and so differs in scale from humans, where BMI is calculated using standing height. Dashed gray line at BMI = 41 shows the cut-off for the analysis in which only males with BMI > 41 were retained for modeling Δ_age_.

**Supplementary Figure 8.**
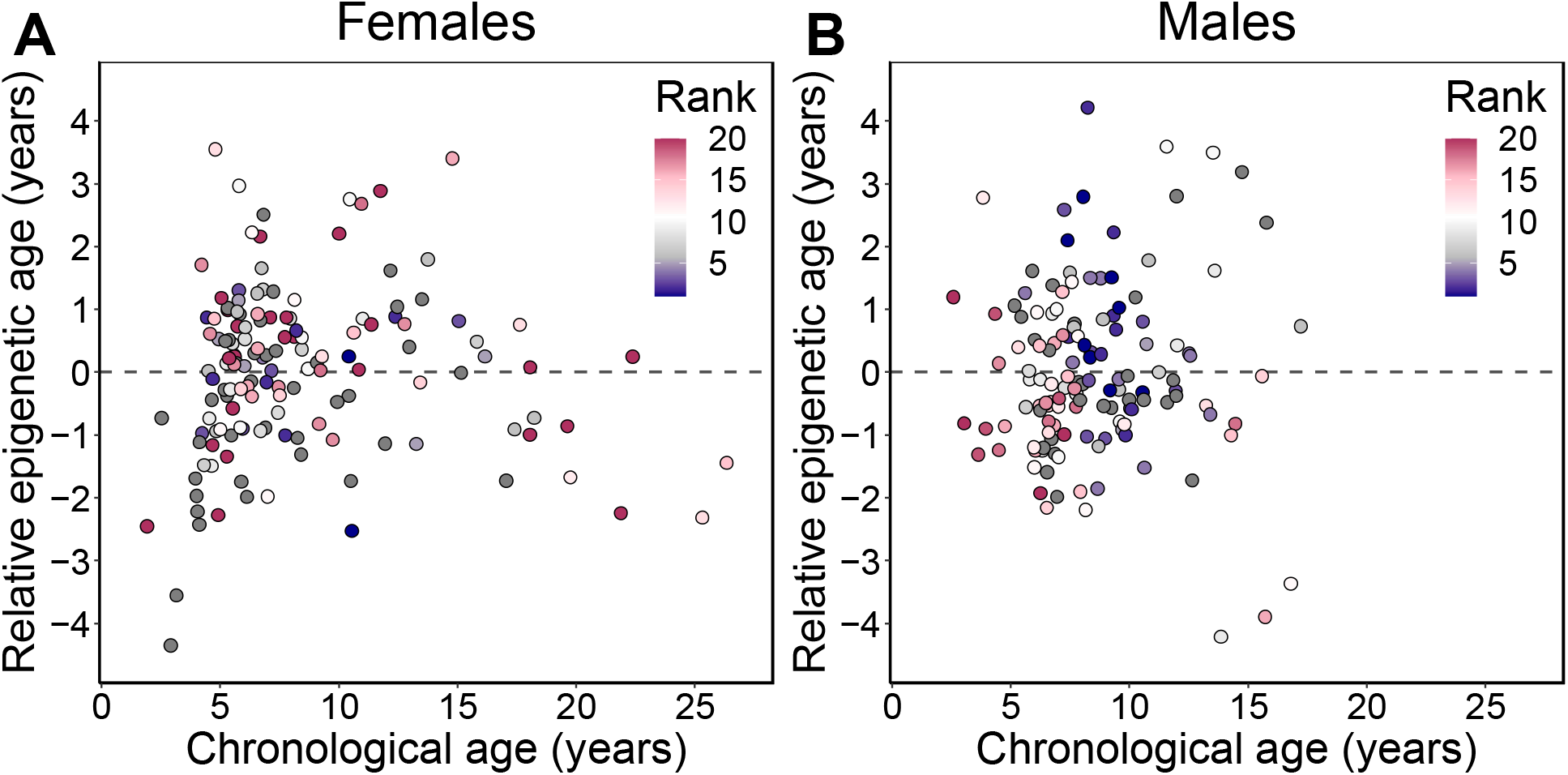
Relative epigenetic age versus chronological age. Each circle represents a baboon, colored by the animal’s dominance rank at the time of sampling. The y-axis shows relative epigenetic age, a measure of epigenetic aging similar to Δ_age_ that is based on the sample-specific residuals from the relationship between predicted age and true chronological age. Positive (negative) values correspond to predicted ages that are older (younger) than expected for that chronological age. Dominance rank is measured using ordinal values, such that smaller values indicate higher rank.

**Supplementary Figure 9.**
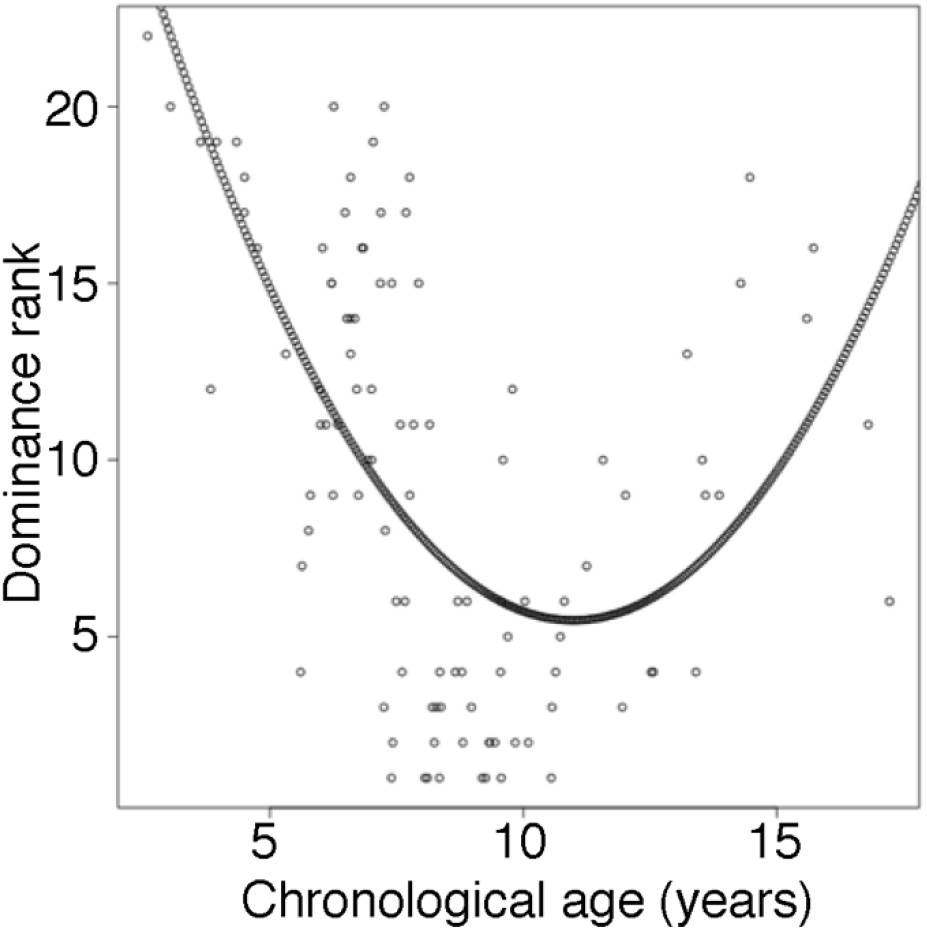
Male dominance rank versus chronological age. Each circle represents a male baboon at the time of sampling. Nearly all males in the top four rank positions are between ages 7 and 12 years (but not all 7 – 12 year olds are also high-ranking: range of rank positions = 1 – 20), whereas both young and old males tend to be lower-ranking. The quadratic curve represents the model with dominance rank as the response variable and age and age^2^ as the predictor variables. Rank-for-age was defined as the residuals of this model.

## References

1 López-Otín, C., Blasco, M. A., Partridge, L., Serrano, M. & Kroemer, G. The hallmarks of aging. Cell 153, 1194–1217 (2013).

2 Jones, O. R. et al. Diversity of ageing across the tree of life. Nature 505, 169 (2014).

3 Monaghan, P., Charmantier, A., Nussey, D. H. & Ricklefs, R. E. The evolutionary ecology of senescence. Functional Ecology 22, 371–378 (2008).

4 Horvath, S. & Raj, K. DNA methylation-based biomarkers and the epigenetic clock theory of ageing. Nature Reviews Genetics 19, 371 (2018).

5 Hannum, G. et al. Genome-wide methylation profiles reveal quantitative views of human aging rates. Molecular cell 49, 359–367 (2013).

6 Horvath, S. DNA methylation age of human tissues and cell types. Genome biology 14, 3156 (2013).

7 Levine, M. E. et al. An epigenetic biomarker of aging for lifespan and healthspan. Aging (Albany NY) 10, 573 (2018).

8 Declerck, K. & Berghe, W. V. Back to the future: Epigenetic clock plasticity towards healthy aging. Mechanisms of ageing and development 174, 18–29 (2018).

9 Levine, M. E., Lu, A. T., Bennett, D. A. & Horvath, S. Epigenetic age of the pre-frontal cortex is associated with neuritic plaques, amyloid load, and Alzheimer’s disease related cognitive functioning. Aging (Albany NY) 7, 1198 (2015).

10 Marioni, R. E. et al. The epigenetic clock is correlated with physical and cognitive fitness in the Lothian Birth Cohort 1936. International journal of epidemiology 44, 1388–1396 (2015).

11 Horvath, S. et al. Obesity accelerates epigenetic aging of human liver. Proceedings of the National Academy of Sciences 111, 15538–15543 (2014).

12 Jovanovic, T. et al. Exposure to violence accelerates epigenetic aging in children. Scientific reports 7, 8962 (2017).

13 Raffington, L. A. S., Belsky, D. W., Malanchini, M., Tucker-Drob, E. M. & Harden, K. P. Analysis of socioeconomic disadvantage and pace of aging measured in saliva DNA methylation of children and adolescents. bioRxiv (2020).

14 Zannas, A. S. et al. Lifetime stress accelerates epigenetic aging in an urban, African American cohort: relevance of glucocorticoid signaling. Genome biology 16, 266 (2015).

15 Maegawa, S. et al. Caloric restriction delays age-related methylation drift. Nature communications 8, 539 (2017).

16 Petkovich, D. A. et al. Using DNA methylation profiling to evaluate biological age and longevity interventions. Cell metabolism 25, 954–960. e956 (2017).

17 Stubbs, T. M. et al. Multi-tissue DNA methylation age predictor in mouse. Genome biology 18, 68 (2017).

18 De Paoli-Iseppi, R. et al. Age estimation in a long-lived seabird (Ardenna tenuirostris) using DNA methylation-based biomarkers. Molecular ecology resources (2018).

19 Polanowski, A. M., Robbins, J., Chandler, D. & Jarman, S. N. Epigenetic estimation of age in humpback whales. Molecular ecology resources 14, 976–987 (2014).

20 Thompson, M. J. An epigenetic aging clock for dogs and wolves. Aging (Albany NY) 9, 1055 (2017).

21 Wright, P. G. et al. Application of a novel molecular method to age free-living wild Bechstein’s bats. Molecular ecology resources 18, 1374–1380 (2018).

22 Alberts, S. C. & Altmann, J. in Long-term field studies of primates 261–287 (Springer, 2012).

23 Lea, A. J., Altmann, J., Alberts, S. C. & Tung, J. Resource base influences genome-wide DNA methylation levels in wild baboons (Papio cynocephalus). Molecular ecology 25, 1681–1696 (2016).

24 Colchero, F. et al. The emergence of longevous populations. Proceedings of the National Academy of Sciences 113, E7681–E7690 (2016).

25 Lea, A. J., Tung, J. & Zhou, X. A flexible, efficient binomial mixed model for identifying differential DNA methylation in bisulfite sequencing data. PLoS genetics 11, e1005650 (2015).

26 Lea, A. J. et al. Genome-wide quantification of the effects of DNA methylation on human gene regulation. eLife 7, e37513 (2018).

27 Lea, A. J. et al. Dominance rank-associated gene expression is widespread, sexspecific, and a precursor to high social status in wild male baboons. Proceedings of the National Academy of Sciences 115, E12163–E12171 (2018).

28 Altmann, J., Gesquiere, L, Galbany, J., Onyango, P. O. & Alberts, S. C. Life history context of reproductive aging in a wild primate model. Annals of the New York Academy of Sciences 1204, 127–138 (2010).

29 Jayashankar, L., Brasky, K. M., Ward, J. A. & Attanasio, R. Lymphocyte modulation in a baboon model of immunosenescence. Clin. Diagn. Lab. Immunol. 10, 870–875 (2003).

30 Galbany, J., Altmann, J., Pérez-Pérez, A. & Alberts, S. C. Age and individual foraging behavior predict tooth wear in Amboseli baboons. American Journal of Physical Anthropology 144, 51–59 (2011).

31 Alberts, S. C. & Altmann, J. Balancing costs and opportunities: dispersal in male baboons. The American Naturalist 145, 279–306 (1995).

32 Alberts, S. C. & Altmann, J. Preparation and activation: determinants of age at reproductive maturity in male baboons. Behavioral Ecology and Sociobiology 36, 397–406 (1995).

33 Alberts, S. C., Watts, H. E. & Altmann, J. Queuing and queue-jumping: long-term patterns of reproductive skew in male savannah baboons, Papio cynocephalus. Animal Behaviour 65, 821–840 (2003).

34 Clutton-Brock, T. H. & Isvaran, K. Sex differences in ageing in natural populations of vertebrates. Proceedings of the Royal Society B: Biological Sciences 274, 3097–3104 (2007).

35 Kirkwood, T. B. & Rose, M. R. Evolution of senescence: late survival sacrificed for reproduction. Philosophical Transactions of the Royal Society of London. Series B: Biological Sciences 332, 15–24 (1991).

36 Williams, G. C. Pleiotropy, natural selection, and the evolution of senescence. evolution, 398–411 (1957).

37 Ryan, J., Wrigglesworth, J., Loong, J., Fransquet, P. D. & Woods, R. L. A systematic review and meta-analysis of environmental, lifestyle and health factors associated with DNA methylation age. The journals of gerontology. Series A, Biological sciences and medical sciences (2019).

38 Tung, J., Archie, E. A., Altmann, J. & Alberts, S. C. Cumulative early life adversity predicts longevity in wild baboons. Nature communications 7, 11181 (2016).

39 Zippie, M. N., Archie, E. A., Tung, J., Altmann, J. & Alberts, S. C. Intergenerational effects of early adversity on survival in wild baboons. eLife 8, e47433 (2019).

40 Archie, E. A., Tung, J., Clark, M., Altmann, J. & Alberts, S. C. Social affiliation matters: both same-sex and opposite-sex relationships predict survival in wild female baboons. Proceedings of the Royal Society B: Biological Sciences 281, 20141261 (2014).

41 Campos, F. A., Villavicencio, F., Archie, E. A., Colchero, F. & Alberts, S. C. Social bonds, social status, and survival in wild baboons: A tale of two sexes. Philosophical Transactions of the Royal Society B: Biological Sciences In press (2020).

42 Holt-Lunstad, J., Smith, T. B. & Layton, J. B. Social relationships and mortality risk: a meta-analytic review. PLoS Med 7, e1000316 (2010).

43 Snyder-Mackler, N. et al. Social determinants of health and survival in humans and other animals. Science 368 (2020).

44 Alberts, S. C., Buchan, J. C. & Altmann, J. Sexual selection in wild baboons: from mating opportunities to paternity success. Animal Behaviour 72, 1177–1196 (2006).

45 Gesquiere, L. R., Altmann, J., Archie, E. A. & Alberts, S. C. Interbirth intervals in wild baboons: Environmental predictors and hormonal correlates. American journal of physical anthropology 166, 107–126 (2018).

46 Lea, A. J., Altmann, J., Alberts, S. C. & Tung, J. Developmental constraints in a wild primate. The American Naturalist 185, 809–821 (2015).

47 Altmann, J., Schoeller, D., Altmann, S. A., Muruthi, P. & Sapolsky, R. M. Body size and fatness of free-living baboons reflect food availability and activity levels. American journal of Primatology 30, 149–161 (1993).

48 Horvath, S. et al. Epigenetic clock and methylation studies in the rhesus macaque. bioRxiv (2020).

49 Engebretsen, S. & Bohlin, J. Statistical predictions with glmnet. Clinical epigenetics 11, 1–3 (2019).

50 Liu, Z. et al. The role of epigenetic aging in education and racial/ethnic mortality disparities among older US Women. Psychoneuroendocrinology 104, 18–24 (2019).

51 Shalev, I. & Belsky, J. Early-life stress and reproductive cost: A two-hit developmental model of accelerated aging? Medical Hypotheses 90, 41–47 (2016).

52 Brody, G. H., Miller, G. E., Yu, T., Beach, S. R. & Chen, E. Supportive family environments ameliorate the link between racial discrimination and epigenetic aging: A replication across two longitudinal cohorts. Psychological Science 27, 530–541 (2016).

53 Brody, G. H., Yu, T., Chen, E., Beach, S. R. & Miller, G. E. Family-centered prevention ameliorates the longitudinal association between risky family processes and epigenetic aging, journal of child psychology and psychiatry 57, 566–574 (2016).

54 Davis, E. et al. Accelerated DNA methylation age in adolescent girls: associations with elevated diurnal cortisol and reduced hippocampal volume. Translational psychiatry 7, e1223 (2017).

55 Marini, S. et al. Predicting cellular aging following exposure to adversity: Does accumulation, recency, or developmental timing of exposure matter? BioRxiv, 355743 (2018).

56 Sumner, J. A., Colich, N. L., Uddin, M., Armstrong, D. & McLaughlin, K. A. Early experiences of threat, but not deprivation, are associated with accelerated biological aging in children and adolescents. Biological psychiatry 85, 268–278 (2019).

57 Austin, M. K. et al. Early-life socioeconomic disadvantage, not current, predicts accelerated epigenetic aging of monocytes. Psychoneuroendocrinology 97, 131–134 (2018).

58 Boks, M. P. et al. Longitudinal changes of telomere length and epigenetic age related to traumatic stress and post-traumatic stress disorder. Psychoneuroendocrinology 51, 506–512 (2015).

59 Lawn, R. B. et al. Psychosocial adversity and socioeconomic position during childhood and epigenetic age: analysis of two prospective cohort studies. Human molecular genetics 27, 1301–1308 (2018).

60 Simons, R. L. et al. Economic hardship and biological weathering: the epigenetics of aging in a US sample of black women. Social Science & Medicine 150, 192–200 (2016).

61 Wolf, E. J. et al. Traumatic stress and accelerated DNA methylation age: a metaanalysis. Psychoneuroendocrinology 92, 123–134 (2018).

62 Aristizabal, M. J. et al. Biological embedding of experience: A primer on epigenetics. Proceedings of the National Academy of Sciences, 201820838 (2019).

63 Hertzman, C. Putting the concept of biological embedding in historical perspective. Proceedings of the National Academy of Sciences 109, 17160–17167 (2012).

64 Ben-Shlomo, Y. & Kuh, D. (Oxford University Press, 2002).

65 Shanahan, L., Copeland, W. E., Costello, E. J. & Angold, A. Child-, adolescent-and young adult-onset depressions: differential risk factors in development? Psychological medicine 41, 2265–2274 (2011).

66 Shanahan, M. J. & Hofer, S. M. in Handbook of aging and the social sciences 135–147 (Elsevier, 2011).

67 Belsky, D. W. et al. Quantification of the pace of biological aging in humans through a blood test, the DunedinPoAm DNA methylation algorithm. Elife 9 (2020).

68 Gesquiere, L. R. et al. Life at the top: rank and stress in wild male baboons. Science 333, 357–360 (2011).

69 Hausfater, G., Altmann, J. & Altmann, S. Long-term consistency of dominance relations among female baboons (Papio cynocephalus). Science 217, 752–755 (1982).

70 Levy, E. J. et al. Higher dominance rank is associated with lower glucocorticoids in wild female baboons: A rank metric comparison. Hormones and Behavior In press.

71 Simons, N. D. & Tung, J. Social status and gene regulation: conservation and context dependence in primates. Trends in cognitive sciences 23, 722–725 (2019).

72 Clutton-Brock, T. et al. Intrasexual competition and sexual selection in cooperative mammals. Nature 444, 1065–1068 (2006).

73 Clutton-Brock, T. H. & Huchard, E. Social competition and selection in males and females. Philosophical Transactions of the Royal Society B: Biological Sciences 368, 20130074 (2013).

74 Emery Thompson, M. & Georgiev, A. V. The high price of success: costs of mating effort in male primates. International Journal of Primatology 35, 609–627 (2014).

75 Alberts, S. C. & Altmann, J. Immigration and hybridization patterns of yellow and anubis baboons in and around Amboseli, Kenya. American Journal of Primatology: Official Journal of the American Society of Primatologists 53, 139–154 (2001).

76 Tung, J., Charpentier, M. J., Garfield, D. A., Altmann, J. & Alberts, S. C. Genetic evidence reveals temporal change in hybridization patterns in a wild baboon population. Molecular Ecology 17, 1998–2011 (2008).

77 Altmann, J. et al. Behavior predicts genes structure in a wild primate group. Proceedings of the National Academy of Sciences 93, 5797–5801 (1996).

78 Tung, J., Zhou, X., Alberts, S. C., Stephens, M. & Gilad, Y. The genetic architecture of gene expression levels in wild baboons. Elife 4, e04729 (2015).

79 Meissner, A. et al. Reduced representation bisulfite sequencing for comparative high-resolution DNA methylation analysis. Nucleic acids research 33, 5868–5877 (2005).

80 Boyle, P. et al. Gel-free multiplexed reduced representation bisulfite sequencing for large-scale DNA methylation profiling. Genome biology 13, R92 (2012).

81 Krueger, F. Trim Galore: a wrapper tool around Cutadapt and FastQC to consistently apply quality and adapter trimming to FastQ files, with some extra functionality for Mspl-digested RRBS-type (Reduced Representation Bisufite-Seq) libraries. URL http://www.bioinformatics.babraham.ac.uk/projects/trim_galore/. (2012).

82 Xi, Y. & Li, W. BSMAP: whole genome bisulfite sequence MAPping program. BMC bioinformatics 10, 232 (2009).

83 Hastie, T., Tibshirani, R, Narasimhan, B. & Chu, G. impute: Imputation for microarray data. Bioinformatics 17, 520–525 (2001).

84 Friedman, J., Hastie, T. & Tibshirani, R. glmnet: Lasso and elastic-net regularized generalized linear models. R package version 1 (2009).

85 Friedman, J., Hastie, T. & Tibshirani, R. Regularization paths for generalized linear models via coordinate descent. Journal of statistical software 33, 1 (2010).

86 Vilgalys, T. P., Rogers, J., Jolly, C. J., Mukherjee, S. & Tung, J. Evolution of DNA methylation in Papio baboons. Molecular biology and evolution 36, 527–540 (2018).

87 Irizarry, R. A. et al. The human colon cancer methylome shows similar hypo-and hypermethylation at conserved tissue-specific CpG island shores. Nature genetics 41, 178 (2009).

88 Consortium, E. P. An integrated encyclopedia of DNA elements in the human genome. Nature 489, 57 (2012).

89 Snyder-Mackler, N. et al. Social status alters immune regulation and response to infection in macaques. Science 354, 1041–1045 (2016).

90 Muggeo, V. M. & Muggeo, M. V. M. Package ‘segmented’. Biometrika 58, 516 (2017).

91 Silk, J. B. et al. Strong and consistent social bonds enhance the longevity of female baboons. Current Biology 20, 1359–1361 (2010).

92 Hausfater, G. Dominance and reproduction in Baboons (Papio cynocephalus). Contributions to primatology 7, 1–150 (1975).

93 Archie, E. A., Altmann, J. & Alberts, S. C. Costs of reproduction in a long-lived female primate: injury risk and wound healing. Behavioral ecology and sociobiology 68, 1183–1193 (2014).

94 Levy, E. J. et al. Comparing proportional and ordinal dominance ranks reveals multiple competitive landscapes in an animal society. bioRxiv (2020).

95 Beehner, J. C., Onderdonk, D. A., Alberts, S. C. & Altmann, J. The ecology of conception and pregnancy failure in wild baboons. Behavioral Ecology 17, 741–750 (2006).

96 Altmann, J. & Alberts, S. C. in Offspring: The Biodemography of Fertility and Family Behavior (eds Kenneth W Wachter & Rodolfo A Bulatao] (The National Academies Press, 2003).

97 Altmann, J., Hausfater, G. & Altmann, S. A. in Reproductive Success: Studies of Individual Variation in Contrasting Breeding Systems (ed Timothy Hugh Clutton-Brock) (The University of Chicago Press, 1988).

98 Charpentier, M., Tung, J., Altmann, J. & Alberts, S. Age at maturity in wild baboons: genetic, environmental and demographic influences. Molecular Ecology 17, 2026–2040 (2008).

99 Altmann, J., Altmann, S. A. & Hausfater, G. Primate infant’s effects on mother’s future reproduction. Science 201, 1028–1030 (1978).

100 Altmann, J. & Alberts, S. C. Variability in reproductive success viewed from a life-history perspective in baboons. American Journal of Human Biology 15, 401–409 (2003).

101 Lea, A. J., Learn, N. H., Theus, M. J., Altmann, J. & Alberts, S. C. Complex sources of variance in female dominance rank in a nepotistic society. Animal behaviour 94, 87–99 (2014).

102 Wang, J. COANCESTRY: a program for simulating, estimating and analysing relatedness and inbreeding coefficients. Molecular ecology resources 11, 141–145 (2011).

